# Downregulation of stromal syntenin sustains AML development

**DOI:** 10.1101/2023.02.15.527799

**Authors:** Raphael Leblanc, Rania Ghossoub, Armelle Goubard, Rémy Castellano, Joanna Fares, Luc Camoin, Stephane Audebert, Marielle Balzano, Berna Bou-Tayeh, Cyril Fauriat, Norbert Vey, Jean-Paul Borg, Yves Collette, Michel Aurrand-Lions, Guido David, Pascale Zimmermann

## Abstract

The crosstalk between cancer and stromal cells plays a critical role in tumor progression. Syntenin is a small scaffold protein involved in the regulation of intercellular communication that is emerging as a target for cancer therapy. Here, we show that certain aggressive forms of acute myeloid leukemia (AML) reduce the expression of syntenin in bone marrow stromal cells (BMSC), stromal syntenin deficiency, in turn, generating a pro-tumoral microenvironment. From serial transplantations in mice and co-culture experiments, we conclude that syntenin-deficient BMSC stimulate AML aggressiveness by promoting AML cell survival and protein synthesis. This pro-tumoral activity is supported by increased expression of endoglin, a classical marker of BMSC, which *in trans* stimulates AML translational activity. In short, our study reveals a vicious signaling loop potentially at the heart of AML-stroma crosstalk and unsuspected tumor-suppressive effects of syntenin that need to be considered during systemic targeting of syntenin in cancer therapy.

## Introduction

Acute myeloid leukemia (AML) is a very heterogeneous malignancy that accounts for over 80% of all acute leukemias in adults, with a five-year overall survival rate below 30% (1). This disease is characterized by the clonal expansion of immature myeloblasts, originating from leukemic stem cells (LSC) which are niched within the bone marrow (BM) (2). The crosstalk between AML cells and surrounding bone marrow stromal cells (BMSC) is essential to sustain tumor progression. Indeed, the leukemic blasts progressively hijack the BMSC to disrupt normal hematopoiesis and promote their own growth and survival (3–6). BMSC-mediated protection of leukemic cells relies on leuko-stromal interactions mediated by ligand-receptor pairs involving adhesion molecules, cytokines, chemokines, growth factors and metabolites. Extracellular vesicles (including exosomes) and mitochondrial transfers also emerged as important vehicles for this cellular cross-talk (7, 8). Identifying specific mechanisms that coordinate the AML-stroma crosstalk can potentially help refining diagnostics and anti-cancer therapies.

Intriguingly, transcriptomic analysis of BMSC isolated from mice inoculated with different subtypes of AML has revealed a downregulation of a protein called syntenin (9). In prior work we identified syntenin as a small PDZ scaffold protein operating in membrane trafficking. It acts as a rate limiting factor in cell-to-cell communication, supporting both *in cis* and *in trans* signaling. Indeed, syntenin works at the cross-section of pathways that mediate the cell surface recycling of endocytosed receptor complexes (10, 11) versus their exosomal secretion after budding into multivesicular bodies (12–17). Syntenin directly binds to multiple transmembrane proteins by virtue of its PDZ domains (15). For instance, by interacting with syndecans, syntenin controls the recycling and exosomal secretion of FGF signals (14, 16), known drivers of cancer cell proliferation and survival (18). Syntenin-syndecan also drives non-cell autonomous SRC-oncogenic effects mediated by exosomes (19). Currently, syntenin gain-of-function is strongly associated with pro-tumoral effects and has been proposed as a therapeutic target (20). Several inhibitors have already been developed for that purpose and tested in preclinical models (21–23). Yet, whether and how stromal syntenin might influence tumoral cell behavior, in particular in AML, is still unknown.

Here we investigate the consequence of stromal syntenin downregulation for AML progression. Using both *in vivo* and *in vitro* approaches and various models of AML, we found that lack of syntenin in the tumor environment, in particular in BMSC, ultimately promotes AML aggressiveness. Exploring the molecular mechanisms at work, both in AML blasts and BMSC, we identified a signaling loop potentially at the heart of stroma-induced tumor survival. This vicious loop is fed by imbalanced stromal endoglin expression and localization, sustained AML EEF1A2/AKT/RPS6 signaling and possibly miR-155 transfer from AML to BMSC.

## Results

### Aggressive AML cells downregulate stromal syntenin expression

By screening a publicly available database (GSE97194) previously established by Lokesh Battula et *al* (9), we found that syntenin expression is downregulated in BMSC isolated from mice with leukemia (**Fig. 1A & Supplementary Table S2**). In this study, BMSC from C57Bl/6 mice transplanted with murine AML cells harboring different genetic alterations (MLL-ENL, MLL-ENL + FLT3-ITD and AML1-ETO9a) were isolated by FACS and their gene expression profiles were established by microarray analysis (9). The selected threshold revealed 241 upregulated (blue dots) and 187 downregulated (orange dots) genes in BMSC stromal cells isolated from mice with leukemia (AML-BMSC) compared to normal BMSC, isolated from non-transplanted animals (**Fig. 1A**). Among those was syntenin (*Sdcbp* gene), whose average expression in BMSC, across various AML genotypes, was 3.2-fold downregulated, compared with control BMSC (**Fig. 1A**). Noteworthy, compared to other AML subtypes, AML cells bearing MLL-ENL + FLT3-ITD mutations seem to be more prone to reduce syntenin expression in BMSC (**Fig. 1B**). Interestingly, the AML FLT3-ITD subtype was shown to overexpress miR-155, an oncogene known to accelerate AML progression (24, 25). Moreover, recent ‘omic’ studies revealed that this miRNA directly downregulates syntenin expression (26–28). To validate that stromal syntenin is indeed a miR-155 target, we first transfected human miR-155 in the human stromal cell line HS5. As expected, we found that miR-155 can reduce syntenin mRNA and protein, by 80% and 60% respectively (**Fig. 1C & 1D**). Of note, when co-transfecting HS5 cells with miR-155 and its inhibitor (anti-miR-155), syntenin expression was no longer affected, illustrating the specificity of the effects (**Fig. 1D**). To test whether human AML cells overexpressing miR-155 can act *in trans* to downregulate stromal syntenin, we co-cultured FLT3-ITD (miR-155^high^) MOLM-14 and AML1-ETO9a (miR-155^low^) U937 cells with HS5 cells (24, 25). After 24h of co-culture, stromal syntenin levels remained unaffected in the presence U937 (miR-155^low^) cells, while MOLM-14 (miR-155^high^) markedly decreased stromal syntenin expression (**Fig. 1E**). Transfecting the stromal cells with miR-155-inhibitor prevented the loss of stromal syntenin induced by MOLM-14 (miR-155^high^) cells, confirming the downregulation of syntenin is indeed mediated by miR-155 (**Fig. 1E**). Altogether, these results indicate that syntenin expression in the microenvironment, in particular in the BMSC, can be modulated by AML cells and that stromal syntenin might be down-regulated by AML expressing high levels of miR-155.

**Figure 1.**
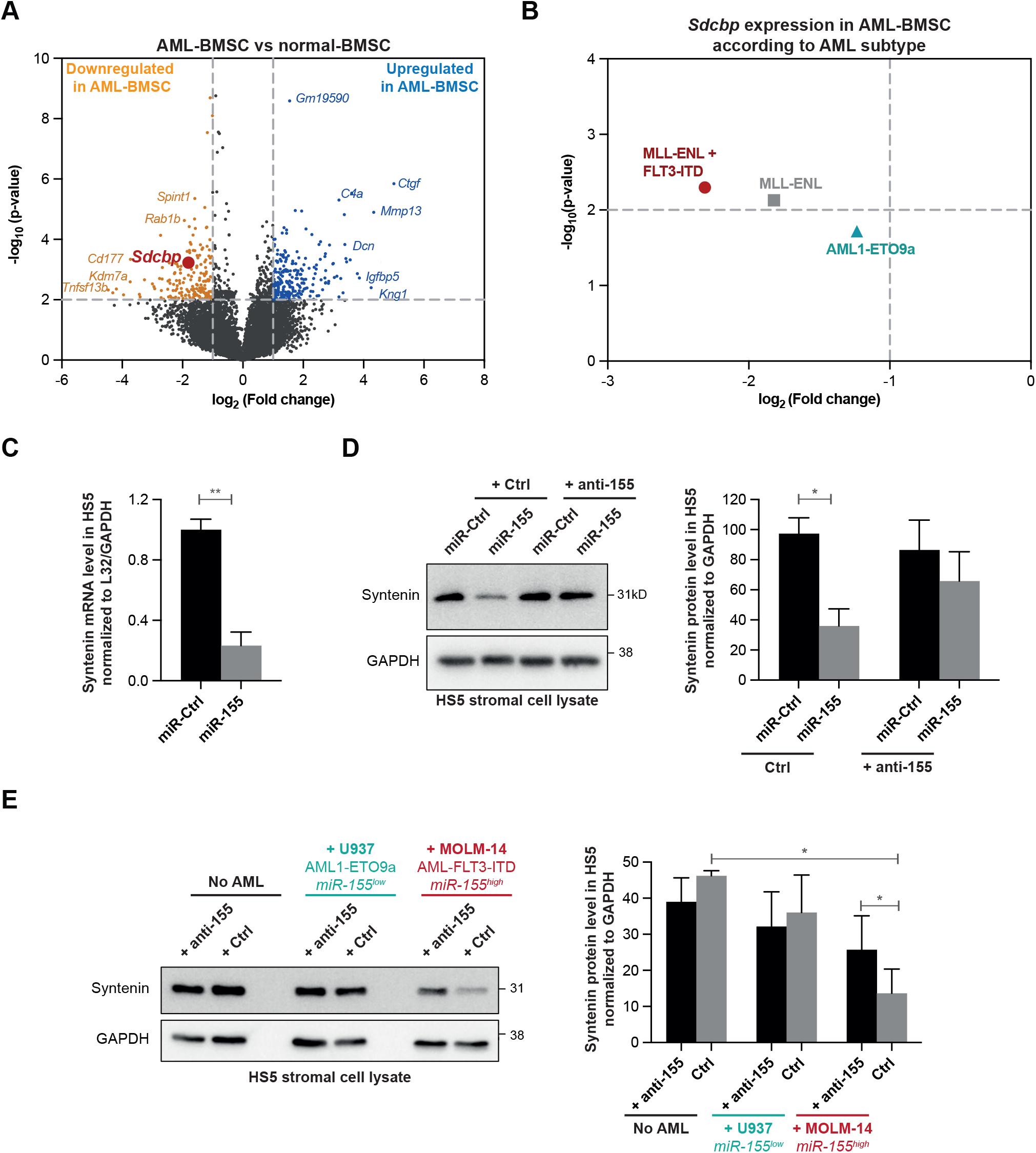
Aggressive human FLT3-ITD^+^ AML downregulate stromal syntenin expression through miR-155. **(A)** The volcano plot was generated from the publicly available database (GSE97194). It illustrates the changes in gene expression, measured by microarray, with log2 fold change (x-axis) and -log10(p-value) (y-axis), in BMSC isolated from C57BL/6 mice inoculated with AML (AML-BMSC) compared to BMSC isolated from control animals (normal-BMSC). Genes, down-or upregulated in AML-BMSC compared normal BMSC, were selected based on a p-value P<0.01 and a difference >2 (see **Supplementary Table S2** for raw data) **(B)** Dot plot representing *Sdcbp* expression in AML-BMSC, compared to normal BMSC, according to the AML subtypes inoculated in the animals (MLL-ENL + FLT3-ITD, MLL-ENL and AML1-ETO9a). The selected threshold is based on a p-value P<0.01 and a difference >2 (see **Supplementary Table S3** for raw data) **(C)** Real-time PCR analysis of syntenin mRNA expression in HS5 transfected with hsa-miR-155-5p (miR-155; 30nM) or control (miR-Ctrl). Values were normalized to housekeeping L32/GAPDH genes. Data represent the mean ± SEM of 3 independent experiments performed in duplicate. Statistical analysis was performed using student t-test (**P < 0.005). **(D)** Left, Western blots on cell lysates from HS5 cells transfected with hsa-miR-155-5p (miR-155) or control (miR-Ctrl), in presence or absence of miR-155-inhibitor (anti-155) illustrating the miR-155 dependent loss of syntenin expression. GAPDH is used as loading control. Right, histograms representing mean syntenin signal intensities ± SEM, relative to GADPH, calculated from the analysis of 3 independent experiments. Statistical analysis was performed using the one-way analysis of variance (ANOVA) (*P < 0.05). **(E)** Left, Western blots of total cell lysates from HS5 transfected with miR-155-inhibitor (anti-155) or control, cultured in absence or presence of U937 (AML-ETO, miR-155^low^) or MOLM14 (AML-FLT3-ITD, miR-155^high^) cells for 24 hours illustrating the impact on syntenin signals. GAPDH was used as loading control. Right, histograms representing mean signal intensities ± SEM, relative to loading control, calculated from the analysis of 3 independent experiments. Statistical analysis was performed using the one-way analysis of variance (ANOVA) (*P < 0.05).

### Host-syntenin deficiency enhances AML aggressiveness

To address the role of host-syntenin in AML development, we made use of the murine FLB1 model that nicely recapitulates features of human AML. FLB1 cells overexpress the oncogenes Meis1 and Hoxa9 and are mir-155^low^ (29). This model rapidly produces widespread AML when inoculated into syngeneic C57Bl/6J mice, the frequency of leukemic stem cells remaining stable over successive transplantations (29). Of note, FLB1 cells are CD45.1, allowing unambiguous identification of the leukemic blasts *versu*s host cells when engrafted in CD45.2 mice. At graft 1, loss of host-syntenin had no noticeable effect on FLB1 progression (**Fig. S1A**). In parallel short-term homing assays, lack of host-syntenin had also no significant effect on the recruitment of the leukemic cells to the primary and secondary hematopoietic organs (**Fig. S1A**). As expected, FLB1 growth in wild-type (WT) animals remained stable over successive transplantations (**Fig. 2**). Yet, FLB1 grown in syntenin-knockout (synt-KO) mice significantly gained in aggressiveness from the 3^rd^ transplantation on (**Fig. 1**) invading bone marrow, spleen and escaping in the peripheral blood within two weeks instead of three (**Fig. S1C**). Altogether, our data reveal that AML blasts confronted to a syntenin deficient microenvironment ultimately acquire an aggressive phenotype.

**Figure 2.**
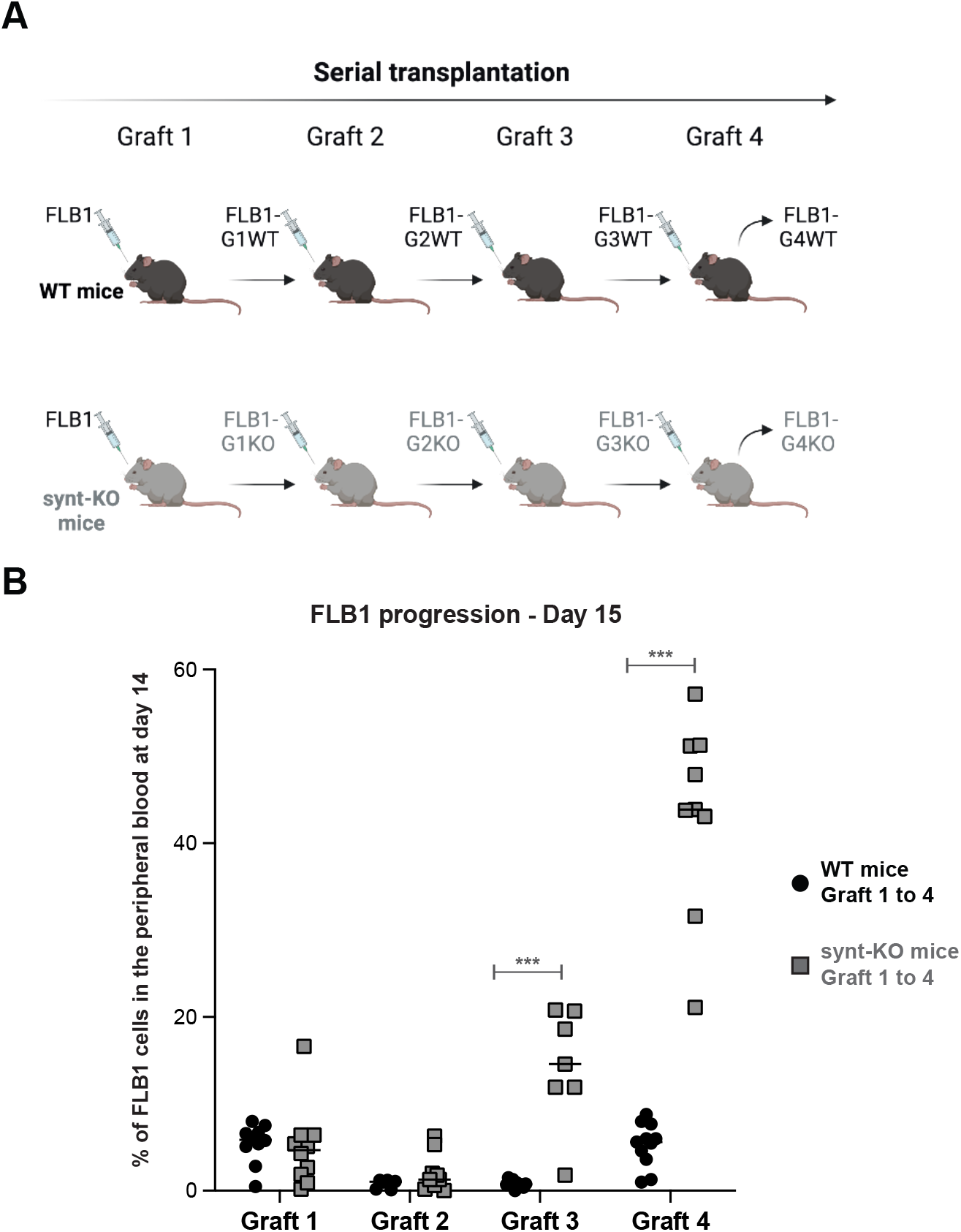
FLB1 blasts gain in aggressiveness upon serial transplantation in a syntenin-deficient host. **(A)** Scheme illustrating the serial transplantation assay. Briefly, WT and syntenin knock-out (synt-KO) mice were injected, in parallel, with FLB1 cells. Upon complete invasion of the bone marrow, FLB1 cells were harvested and re-injected for a new round of transplantation. FLB1 were serially transplanted into WT and synt-KO animals in parallel for 4 rounds (graft 1 to graft 4) **(B)** FACS analysis of blood samples collected from mice on day 14 post injection. Leukemia burden is expressed as percentage of FLB1 (CD45.1^+^) cells relative to total CD45^+^ (CD45.1^+^+CD45.2^+^) in the peripheral blood ± SEM. Statistical analysis was performed using the two-way analysis of variance (ANOVA) (**P < 0.005; ***P < 0.0005). Note the stability of the leukemia burden upon serial transplantation in WT animals. Note the leukemia ‘outbreak’, from graft 3 on, in synt-KO animals.

### AML blasts educated in a syntenin-deficient host display enhanced survival and protein synthesis

To identify the mechanisms supporting this AML aggressiveness, we first assessed classical “hallmarks” associated with leukemia progression. As shown in **figures S2A** to **S2C**, we excluded an altered homing, change in leukemic initiating cell frequencies or cell cycle defect. We then compared the proteomes of FLB1 blasts isolated from WT and synt-KO animals after graft 4 (further referred to as FLB1-G4WT and FLB1-G4KO, respectively). Quantitative protein expression analysis by mass spectrometry allowed the reproducible identification of 2,705 proteins (**Supplementary Table S5**). **Figure 3A** shows a volcano plot of the entire data set, highlighting proteins whose expression were significantly different within FLB1-G4WT and G4KO cells. The selected thresholds revealed 104 up-regulated (blue dots) and 112 down-regulated (orange dots) proteins in FLB1-G4 isolated from synt-KO animals. Bioinformatic evaluation of these changes (i.e. Ingenuity Pathway Analysis) identified ‘Cell Death and Survival’ and ‘Protein Synthesis’ pathways, as the top two molecular and cellular functions that were significantly different (**Fig. 3B**). To validate these observations, we first tested the capacity of FLB1-G4 cells to survive *ex vivo*. FLB1-G4 cells were isolated from bone marrow aspirates by FACS and seeded in complete medium. We found that FLB1-G4KO undergo little apoptosis (**Fig. 3C**). In fact, the propensity of these cells to survive is exacerbated so that they can easily be grown *ex-vivo*, unlike FLB1transplanted in WT animals. We also compared the translational activity of FLB1-G4WT and FLB1-G4KO *in vivo*, in WT and synt-KO animals respectively, using O-propargyl-puromycin labelling methods (**Fig. 3D**) (30). AML blasts evolving in synt-KO mice showed a significant 2-fold increase in total protein synthesis (**Fig. 3D & S2D**). Total protein synthesis in the nonmalignant (CD45.1^-^) host cells, in contrast, was similar in WT and synt-KO mice (**Fig. 3C & S2D**). Assessing classical pathways associated with cell survival and protein translation, we found that FLB1-G4KO cells display higher cellular levels of total AKT, associated with increased levels of AKT phosphorylated on S473 and T450 (**Fig. 3E**). In contrast, FLB1-G4KO show no activation of STAT3, suggesting specific activation of survival pathways (**Fig. S2E**). We additionally tested for the activation of RPS6, a downstream effector of AKT involved in protein synthesis. Interestingly, FLB1-G4KO cells showed a significant increase of the cellular level of RPS6, as well as increased levels of phosphorylated S235-RPS6 (**Fig. 3D**).

**Figure 3.**
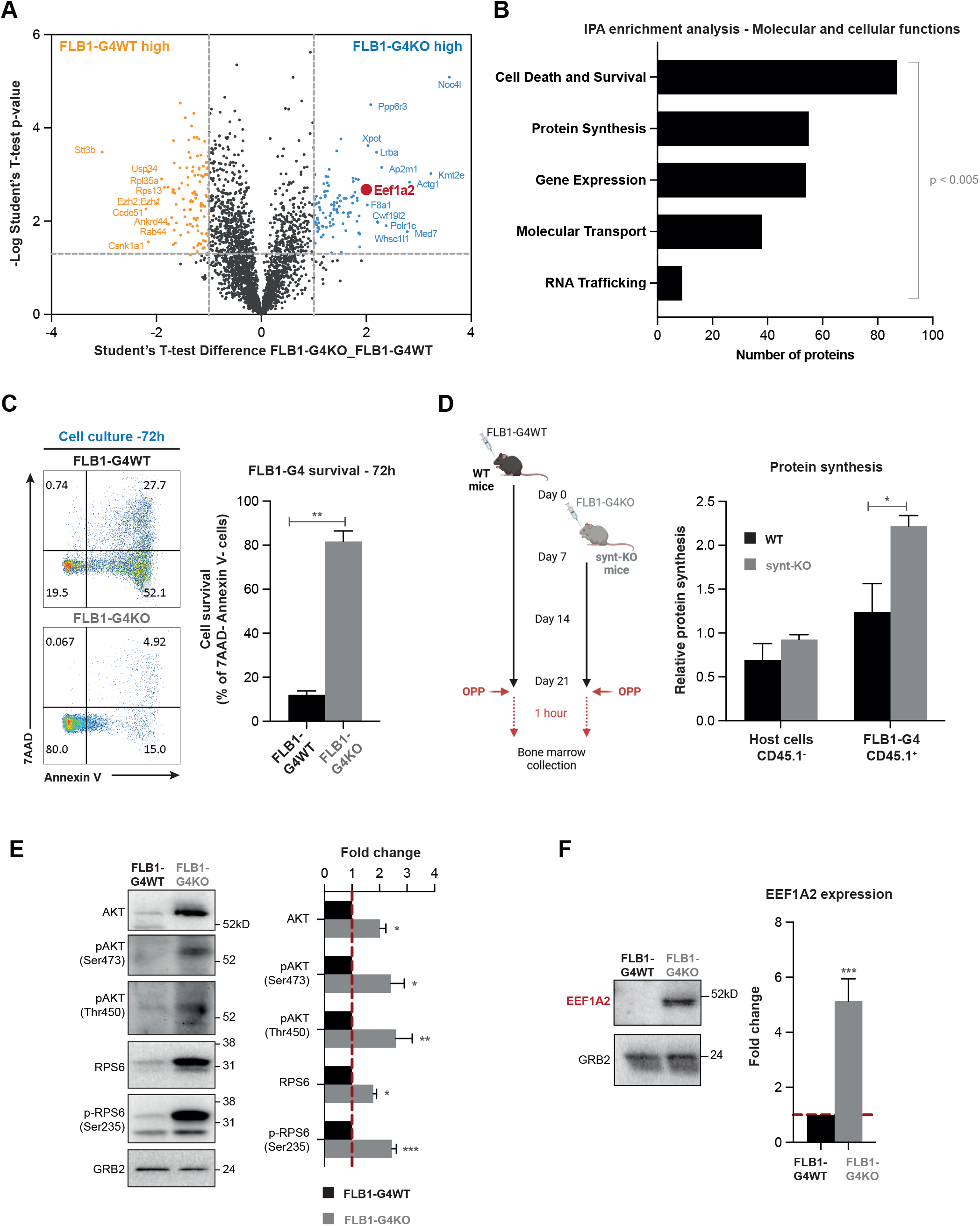
Education by a syntenin-deficient host sustains cell survival and protein synthesis in leukemia blasts. **(A)** Volcano plot illustrating the proteins that are differentially expressed, with log10 levels (x-axis) and –log(p-value) (y-axis), in FLB1-G4KO (aggressive) blasts versus FLB1-G4WT blasts (see **Supplementary Table S5** for raw data). **(B)** Molecular and cellular functions identified using ingenuity pathway analysis (IPA), based on the selected threshold (P<0.005; Difference>1). The bar diagram indicates the most significant molecular and cellular functions upregulated in FLB1 cells transplanted for 4 rounds in Synt-KO mice compared to FLB1 cells transplanted in WT animals. The x-axis indicates the number of molecules involved in the corresponding y-axis function. **(C)** FACS analysis of FLB1 (CD45.1^+^) cells collected after graft 4 from WT (FLB1-G4WT) or synt-KO animals (FLB1-G4KO). Apoptosis was evaluated after 72h of maintenance in complete RPMI media, using double staining with 7AAD/AnnexinV, as illustrated on the representative dot blots (Left panel). Results from 3 independent experiments are expressed as mean value of living (annexinV^-^, 7AAD^-^) FLB1cells ± SEM. Statistical analyses were performed using the nonparametric Mann-Whitney U test (**, P < 0.001) (Right panel). **(D)** Left, scheme of the experiment. FLB1-G4WT cells were injected at day 0 into WT mice, while the inoculation of FLB1-G4KO cells into synt-KO mice was delayed for one week to obtain similar stages of disease progression in the two different hosts at day 21. To address *in vivo* protein synthesis, O-propargyl-puromycin (OPP) was injected intraperitoneally. One hour later, BM was collected and OPP incorporated into nascent polypeptide chains was fluorescently labeled via “Click-it Chemistry”. Right, Graph representing the relative levels of protein synthesis in host cells (CD45.1-) and in FLB1 cells (CD45.1+) relative to the BM cells from OPP-untreated mice ± SEM as measured by FACS and calculated from the analysis of 3 independent mice. Statistical analysis was performed using the nonparametric Mann-Whitney U test (*, P<0.05). **(E)** Left, Western blots of FLB1-G4WT and -G4KO total cell lysates analyzed for different markers, as indicated. Right, histograms representing FLB1-G4KO mean signal intensities ± SEM, relative to signals obtained with FLB1-G4WT lysates, calculated from the analysis of 5 independent mice. Statistical analysis was performed using the nonparametric Mann-Whitney U test (*, P<0.05; **, P<0.001; ***, P < 0.0001). **(F)** Left, Western blot from FLB1-G4WT and -G4KO total cell lysates illustrating EEF1A2 expression. The ubiquitous GRB2 signal was used as loading control. Right, histogram representing FLB1-G4KO EEF1A2 mean signal intensities ± SEM, relative to signals obtained with FLB1-G4WT lysates, calculated from the analysis of 5 independent mice. Statistical analysis was performed using the nonparametric Mann-Whitney U test (***, P < 0.0001).

In an attempt to identify a ‘molecular node’ associated with the activation of these pathways, we carefully analyzed the proteins differentially expressed in FLB1-G4KO cells. Interestingly, we noted the Eukaryotic Translation Elongation Factor 1 Alpha (EEF1A2) ranked among the top up-regulated factors (**Fig. 3A**). EEF1A2 is a translation factor implicated in the delivery of aminoacyl-tRNAs to the A site of the ribosome (31). It is also known as a proto-oncogene, whose expression is linked to anti-apoptotic functions in several types of cancer, including AML, through the activation of AKT/mTORC1/RPS6 signaling (32–34). Western blot confirmed that EEF1A2 was significantly upregulated, nearly 5-fold (**Fig. 3F**). Then, we investigated to what extent gain of EEF1A2 can explain the gain of AML survival and aggressiveness. When treating aggressive FLB1-G4KO cells, *ex vivo*, with increasing concentrations of Metarrestin, an EEF1A2 inhibitor (35), we observed reduced cellular levels of pS473-AKT, pT450-AKT, RPS6 and pS235-RPS6 (**Fig. S3A**). Metarrestin treatment was also associated with a dose-dependent apoptosis of aggressive FLB1-G4KO cells *ex vivo* (**Fig. S3B**). As mice did well tolerate Metarrestin treatment for up to 14 days (**Fig. S3C-D**), we also tested whether such treatment would impact on bone marrow invasion by FLB1-G4KO and FLB1-G4WT. While we observed no effect of the treatment on FLB1-G4WT, we found that Metarrestin had a significant effect on bone marrow invasion by FLB1-G4KO (**Fig. S3E**).

Altogether, our data reveal that a syntenin-deficient environment may ultimately lead to the activation of EEF1A2/AKT/RPS6 molecular pathways in AML cells, stimulating protein synthesis and enhancing cell survival, advantages educating the leukemia to become more aggressive.

### *Ex-vivo* long-term co-culture with syntenin-deficient BMSC suffices to convey an AML aggressive phenotype

BMSC were previously reported to support AML aggressiveness by diverse mechanisms (3, 36, 37). We therefore aimed to clarify whether syntenin-KO BMSC might account for the AML aggressive phenotype. No conditional knockout targeting BMSC is available, as this is a highly heterogenous population. We therefore isolated BMSC from both WT and synt-KO mice and expanded them *in vitro* (**Fig. S4A**). BMSC from both sources appear similar in terms of differentiation potential (**Fig. S4B**). To test whether synt-KO BMSCs would be competent to educate AML *in vitro*, we kept FLB1 cells in culture, in the presence of BMSC, WT or synt-KO, for one month (**Fig. 4A**). Of note, original FLB1 cells were difficult to maintain *ex vivo*, in particular when co-cultured with BMSC WT, and considerable amounts of cells were required to conduct these experiments. After one-month of co-culture, the remaining FLB1 (CD45.1^+^) cells were collected and injected into WT animals. At the moment the disease just became detectable in animals injected with FLB1 maintained on a WT stroma, we noticed an already important invasion of the peripheral blood, the bone marrow and the spleen when FLB1 had been co-cultured with synt-KO BMSC (**Fig. 4A**). This result demonstrates that exposure to synt-KO stromal cells alone suffices to educate AML cells to become more aggressive.

**Figure 4.**
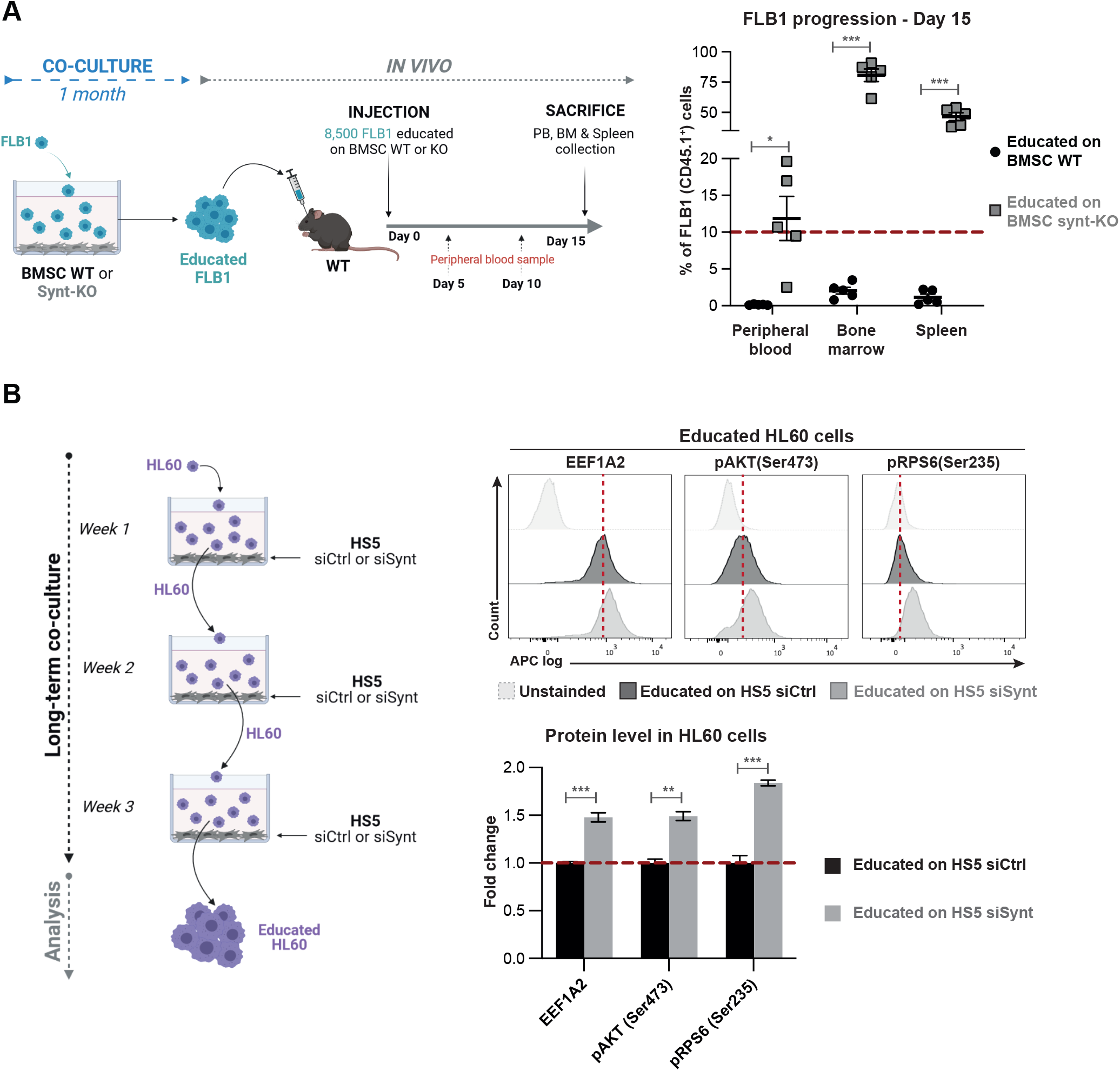
Syntenin-deficient BMSC suffice to educate AML cells for aggressiveness. **(A)** Left, scheme for *ex-vivo* long-term FLB1-BMSC co-culture experiments coupled to *in vivo* leukemia burden assays. FLB1 cells were co-cultured with murine BMSC isolated from WT or synt-KO mice. After one month of co-culture, surviving FLB1 cells, ‘educated’ by murine BMSC (WT or synt-KO), were injected in the retro-orbital vein of WT mice. Leukemia progression was assessed weekly, by FACS analysis as described before. Right, results of FACS analyses showing FLB1 (CD45.1+) cell frequencies measured in PB, BM and the spleen of the different mice in the different groups, 15 days after cell inoculation. Results are expressed as percentage of FLB1 (CD45.1^+^) cells relative to total CD45^+^ cells, and calculated means ± SEM. Statistical analysis was performed using one-way ANOVA test (*, P < 0.05; **, P < 0.005). **(B)** Left, scheme for co-culture experiments with human leukemia and stromal cells. HS5 (stroma) cells were transfected with siRNA targeting syntenin (siSynt) or control (siCtrl). 24h later, human AML HL60 cells were brought in contact with HS5 cells. After one week of co-culture, HL60 cells were collected and re-seeded on freshly transfected HS5 cells. After 3 weeks, HL60 cells ‘educated’ by HS5-siCtrl or HS5-siSynt were collected for FACS analysis. Right, Representative FACS analysis, of EEF1A2, pS473-AKT and pS235-RPS6 protein levels in HL60 cells after 1 month of co-culture with HS5 transfected with siRNA targeting syntenin (siSynt) or control (siCtrl). Histogram representing fold change in mean of fluorescence intensity relative to educated HL60 maintained on HS5 transfected with siCtrl ± SEM calculated from the analysis of 3 independent experiments. Statistical analysis was performed using the two-way analysis of variance (ANOVA) (**P < 0.005; ***P < 0.0005).

We then tested whether a similar phenomenon may be observed with human models. For that, we made use of the stromal cell line HS5 and the AML cell lines HL60 (mutated for Nras and Cdkn2a) and U937 (mutated for p53). HS5 were silenced for syntenin expression and co-cultured with human AML cells for one month (**Fig. 4B & S4C**). To maintain the level of stromal syntenin at its minimum during the whole experiment, stroma-exposed AML cells were collected each week and re-seeded on freshly transfected (silenced) HS5 cells. In agreement with our *in vivo* observations on FLB1 cells, co-culture with syntenin-deficient HS5 stromal cells increased EFF1A2 expression in both the HL60 (**Fig. 4B**) and U937 AML models (**Fig. S4C**). In HL60 cells educated on syntenin-deficient HS5 stromal cells, this increase was associated with higher levels of phosphorylated S473-AKT and S235-RPS6 (**Fig. 4B**). Moreover, we noticed that syntenin-deficiency in HS5 stromal cells markedly stimulates the protein synthesis in HL60 cells (see **Fig. 4E**, siCtrl vs siSynt + SiCtrl). Taken together, these results indicate that a syntenin-deficient stroma may stimulate AML aggressiveness in both mice and humans.

### Syntenin-deficient BMSCs display altered endoglin distribution supporting the AML phenotypic change

To clarify the pro-tumoral mechanisms at work in syntenin-deficient stroma, we compared the mesenchymal patterns of WT and synt-KO murine BMSC. We noticed markedly increased levels of endoglin protein at the surface of synt-KO BMSC, compared to WT (**Fig. 5A**). Consistent with the conservation of the effects between mice and humans, syntenin-deficiency in HS5 stromal cells is also accompanied by an increase of endoglin protein at the cell surface (**Fig. 5B**) and of total cellular endoglin (**Fig. 5C**). Because of the important role of syntenin in exosomal releases, we tested for a potential defect of vesicular endoglin secretion. Comparing small extracellular vesicles (small EVs) purified by differential ultracentrifugation (enriched in exosomes), we found a significant decrease of endoglin levels into the small EVs isolated from syntenin-deficient HS5 cells (**Fig. 5C**). Interestingly, treatment of HS5 with miR-155 mimics the effects of syntenin deficiency, increasing the levels of cell-surface and cellular endoglin (**Fig. S5A & S5B**) and decreasing endoglin levels associated with extracellular vesicles (**Fig. S5B**). Endoglin is a transmembrane glycoprotein modulating TGF*β* signaling, notably through its PDZ interactions (38). As PDZ interactions are known to be promiscuous (39) and as syntenin is a PDZ protein, we investigated whether syntenin might directly bind endoglin. By surface plasmon resonance (SPR) experiments, we found that the PDZ domains of syntenin interact directly with endoglin cytosolic domain, in a concentration-dependent manner (**Fig. 5D**), similarly to GIPC1, a previously described endoglin-PDZ interactor (**Fig. S5C**). The apparent KD values were determined to be 27.1μM and 2.4μM respectively (**Fig. S5D**). In addition, confocal imaging revealed that endogenous endoglin partially colocalizes with endogenous syntenin, at intracellular sites (**Fig. 5E**).

**Figure 5.**
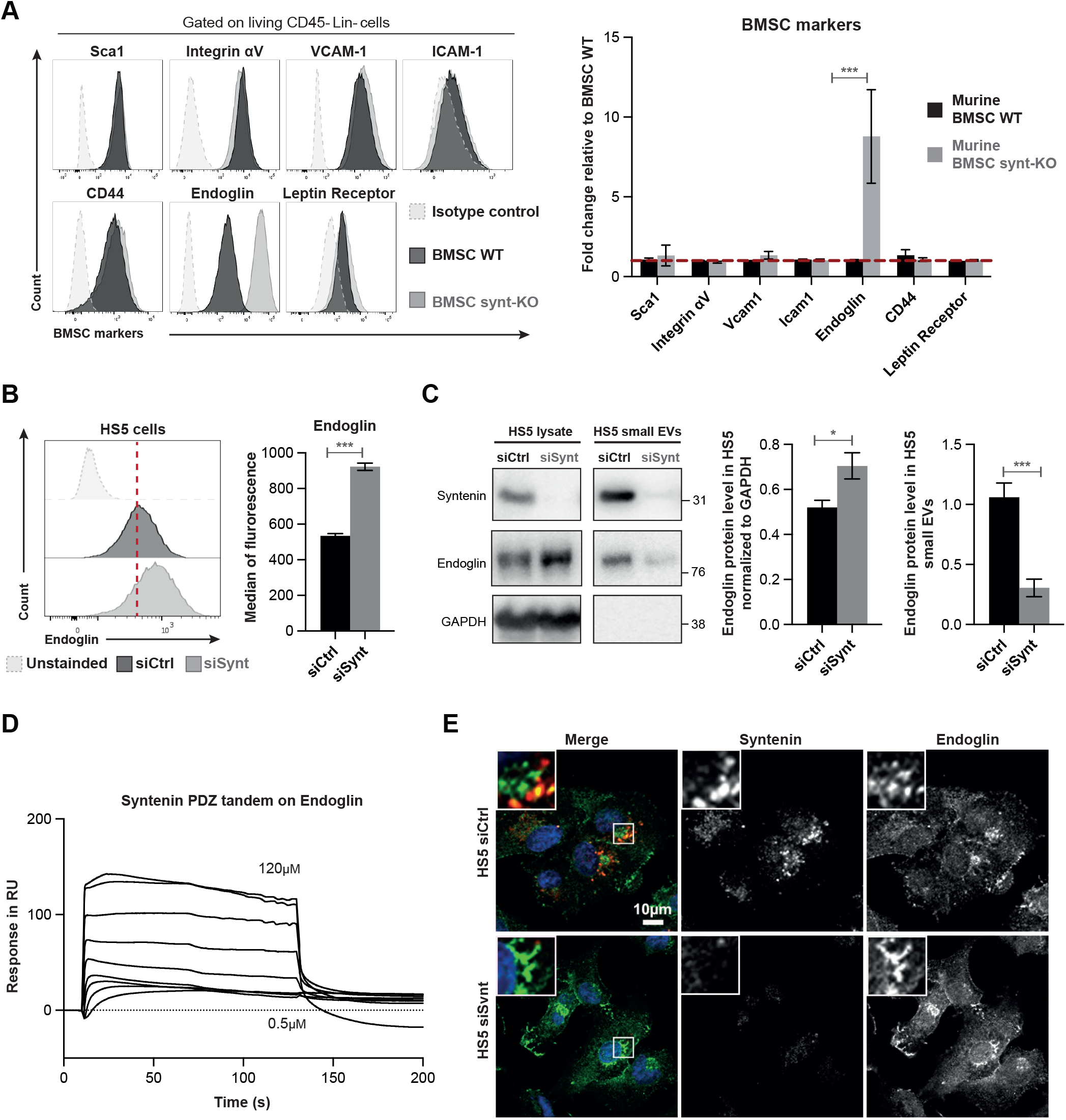
Syntenin-deficiency in BMSC affects the distribution of endoglin. **(A)** Right, FACS-analysis of cell surface BMSC markers (as indicated) in WT (black lane) or Synt-KO (gray lane) murine expanded BMSCs (passage 4). Right, histogram representing the fold change in the median of fluorescence relative to that measured in BMSC WT ± SEM, calculated from the analysis of 5 independent experiments. Statistical analysis was performed using parametric student t-test (***P < 0.0005). **(B)** Left, FACS detection of cell surface expression of endoglin in HS5 cells transfected with siRNA targeting syntenin (siSynt; gray histogram) or control (siCtrl; black histogram). Right, histogram representing the median of fluorescence of endoglin ± SEM. Values were calculated from 3 independent experiments. Statistical analysis was performed using the one-way analysis of variance (ANOVA) (*** P < 0.0005). **(C)** HS5 cells were transfected with siRNA targeting syntenin (siSynt) or control siRNA (siCtrl). After 48h, the media were changed and HS5 transfected cells were cultured in medium containing EV-depleted FCS (10%) for 16h. Conditioned medium was submitted to differential centrifugation and small EVs were pelleted at 100,000g. Upper, total cell lysates and corresponding pelleted extracellular particles (small EVs) were analyzed for indicated markers. Lower, histograms represent the endoglin protein levels in HS5 cell lysate (normalized to GAPDH) and in HS5 small EVs. Values were calculated from 3 independent experiments. Statistical analysis was performed using parametric student t-test (* P < 0.05; ***P < 0.0005). **(D)** Surface plasmon resonance experiment illustrating the direct syntenin-endoglin interaction, tested with recombinant polypeptides. Increasing concentrations of syntenin (comprising only the tandem-PDZ domains + C-terminal domain; 0.5μM to 120μM) were perfused on an immobilized peptide corresponding to the last 25 C-terminal amino acids of wild-type endoglin. **(E)** Representative confocal micrographs showing the steady-state distribution of endogenous syntenin and endogenous endoglin in HS5 cells depleted for syntenin expression (HS5 siSynt) compared to control cells (HS5 siCtrl). In merge, nuclei are stained with DAPI (blue), syntenin is in red, and endoglin is in green.

We finally assessed whether endoglin may be implicated in the acquisition of the aggressive AML phenotype conveyed by a syntenin-deficient stroma. For that, HL60 cells were exposed to HS5 stromal cells silenced for endoglin (**Fig. 6A, S6A & S6B**). We noticed that stromal endoglin deficiency significantly reduces protein synthesis and the phosphorylation of S473-AKT and S235-RPS6 in HL60 (**Fig. 6B & 6C**). Thus, endoglin deficiency has opposite effects compared to syntenin deficiency (**Fig. 4B, Fig. 6B & Fig. 6C**). When silencing both syntenin and endoglin in HS5 cells (**Fig. S6A & S6B**), the level of protein synthesis and the levels of pS473-AKT & pS235-RPS6 in HL60 return to a “baseline level”, similar to the level observed in HL60 co-cultured with HS5 control cells (**Fig. 6B & 6C**). Altogether, these data show that syntenin controls the expression of endoglin in stromal cells, and suggest that an ‘endoglin high’ stroma may be implicated in the acquisition of the aggressive AML phenotype promoted by a syntenin-deficient stroma.

**Figure 6.**
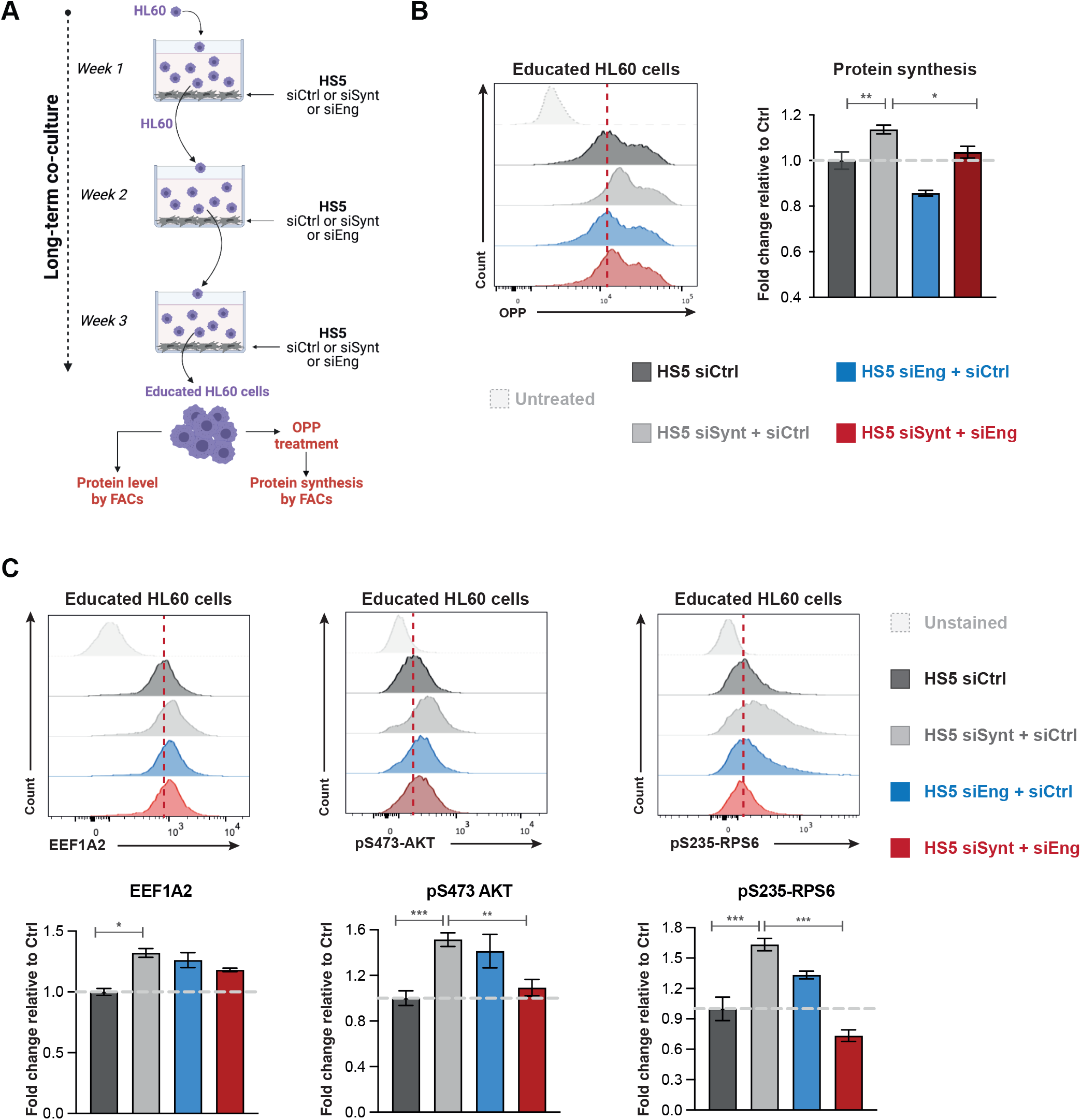
High endoglin expression in BMSC acts *in trans* to regulate AML protein synthesis. **(A)** Scheme illustrating long term co-culture experiments with syntenin- and endoglin-deficient stromal cells, addressing stromal effects on AML protein synthesis. HS5 cells were transfected with siRNA targeting syntenin (siSynt), endoglin (siEng) or control siRNA (siCtrl). 24h later, human AML HL60 cells were co-cultured with these HS5 cells. After one week of co-culture, HL60 cells were collected and re-seeded on freshly transfected HS5 cells. After 3 weeks of co-culture, HL60 AML cells were collected and treated with O-propargyl-puromycin (OPP) to measure protein synthesis or analyzed by FACs for markers. **(B)** Left, FACS detection of OPP incorporation in HL60 educated on HS5 treated as indicated. Untreated refers to HL60 not treated with OPP. Right, graph representing the relative levels of protein synthesis in HL60 cells, normalized to untreated HL60 cells ± SEM. Values were calculated from 4 independent experiments. Statistical analysis was performed using the one-way analysis of variance (ANOVA) (*, P<0.05; **, P<0.005). **(C)** Upper, representative FACS analysis of EEF1A2, pS473-AKT and pS235-RPS6 protein levels in HL60 cells after 1 month of co-culture with HS5 transfected with siRNA targeting syntenin (siSynt), endoglin (siEng), both (siSynt + siEng) or control (siCtrl). Histogram representing the fold change in mean of fluorescence intensity relative to educated HL60 maintained on HS5 transfected with siCtrl ± SEM calculated from the analysis of 3 independent experiments. Statistical analysis was performed using the two-way analysis of variance (ANOVA) (**P < 0.005; ***P < 0.0005).

## Discussion

Using an AML-relevant syngeneic mouse model and *in vitro* human and mouse co-culture assays, we here reveal several unsuspected molecular mechanisms that may govern tumor-stroma communication. Indeed, our study indicates that down-regulation of stromal syntenin might be at the heart of a pro-tumoral vicious signaling loop between the tumor cells and the associated bone marrow stromal cells (**Fig. 7**).

**Figure 7.**
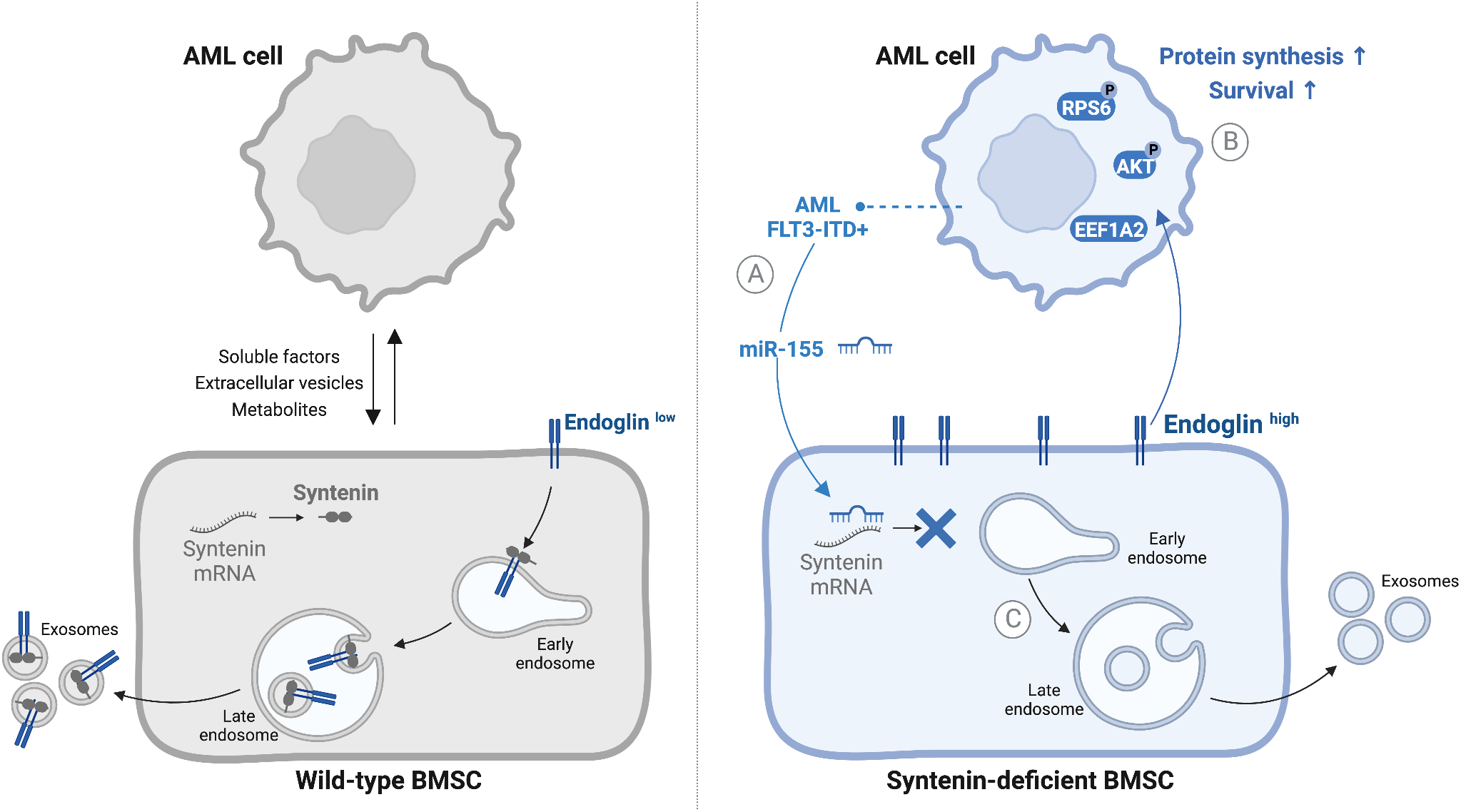
Model recapitulating novel findings presented in this study. We here provide novel insight in the mechanisms at work in tumor-stroma crosstalk. **(A)** Aggressive AML (FLT3-ITD) cells overexpressing miR-155 can suppress syntenin expression in stromal cells. **(B)** AML blasts confronted with a syntenin-deficient stroma acquire a cell survival advantage associated with increased levels of protein synthesis. This relies on the activation of the EEF1A2/AKT/RPS6 pathway. **(C)** Syntenin dysregulation increases the level of stromal endoglin at cell surfaces but decreases the loading of endoglin into exosomes. High stromal endoglin is needed for the *in trans* support of AML translational activity by a syntenin-deficient stroma.

In the context of AML, we found that such signaling loop might be initiated by ‘aggressive’ blasts, in particular AML of the FLT3-ITD^+^ subtype overexpressing miR-155. Indeed, AML that are miR-155 high are able to down-regulate stromal syntenin in co-culture (**Fig. 1E**). A phenomenon that is not observed with AML expressing low levels of miR-155. This is intriguing because the pro-tumoral effect of miR-155 is currently primarily considered to take place via mechanisms occurring in the tumor cells. This holds for AML blasts, and also for glioma, lung, colorectal or breast cancers (40). We hereby provide evidence that the pro-tumoral effect of miR-155 may also depend on effects that occur *in trans*, i.e. via downregulation of stromal syntenin (**Fig. 1E**). The publicly available data on *in vivo* expressions indicate that also other AML subtypes bearing different genetic alterations, in particular MLL-ENL, are capable of reducing syntenin expression in BMSC (**Fig.1B**). This suggests that also other factors might be involved in the downregulation of stromal syntenin expression. miR-7977 might be a relevant candidate since it can be transferred from different subtypes of AML cells to BMSC (41) and syntenin is one of its main targets (42). Therefore, the regulation of stromal-syntenin expression in distinct AML subtypes requires deeper analysis.

The finding that leukemic blasts confronted with a syntenin-KO environment gain in aggressiveness was completely unexpected (**Fig. 2B**). Indeed, up to now, syntenin was unambiguously considered as a valuable target for cancer therapy and several inhibitors have already been developed and tested in preclinical models (20, 22, 23, 43). This is because gain of syntenin expression in tumor cells has been invariably associated with the invasion and the metastatic potential of various solid tumors, including melanoma, glioblastoma, breast, prostate and head/neck squamous cancers (20). Moreover, syntenin-KO mice were previously shown to be refractory to melanoma metastasis due to reduced tumor-supporting inflammation (44). One reason for this apparent controversy may be that the B16 melanoma tumor model used in these (single graft) experiments was only shortly exposed to the syntenin-deficient microenvironment, while our study addressed the effects of long-term exposure to a syntenin-deficient stroma (**Fig. 1B & 3A-B**). Another trivial explanation might be that a syntenin-deficient stroma has different consequences depending on the tumor type. That remaining to be clarified, our study invites to also considerate the long-term effects of anti-syntenin therapy in the context of cancer treatment. It implies that the suitability and efficacy of syntenin as a systemic target in cancer therapy (including the emergence of drug resistance) may ultimately also depend on monitoring and compensating for environmental loss of syntenin activity.

Remarkably, the spectacular gain of survival acquired by AML blasts facing a syntenin-KO environment (**Fig. 3B**) is associated with a 2-fold increase in tumor translational activity (**Fig. 3C**). In the last decade, aberrant translation emerged as an important player in the pathogenesis of cancers (45). The precise molecular mechanism by which stroma is supporting high protein synthesis in the tumor remaining enigmatic, we here show that stromal syntenin-deficiency is inducing a 5-fold increase of the elongation factor EEF1A2 (**Fig. 3A, 3F & 7B**) in AML. In addition to its role in aminoacyl-tRNA delivery to the ribosome, EEF1A2 is known for its pro-oncogenic activity in many tumors, enhancing cancer cell proliferation and inhibiting apoptosis (31). Moreover, EEF1A2 is known to activate the pro-survival AKT signaling pathway (31, 33, 46). This is completely consistent with our observations that AML blasts educated in a syntenin-KO environment display activated AKT/RPS6 signaling and survive better (**Fig. 3E & 4C**). It is thus tempting to propose that EEF1A2 works as a node supporting the pro-tumoral effects elicited by the stromal deficiency of syntenin. In agreement with this contention, we observed that pharmacological inhibition of EEF1A2 by Metarrestin, *in vitro*, drastically induces apoptosis of AML blasts educated by a syntenin-KO stroma (**Fig. S3B**). Moreover, Metarrestin treatment significantly reduces the aggressiveness of blasts educated in a syntenin-KO host but has no effect on AML blasts grown in wild-type mice (**Fig. S3E**).

In an effort to identify cues to mechanisms supporting the stromal syntenin-KO pro-tumoral effects, we found that the TGFβ co-receptor endoglin is highly expressed at cell membranes in syntenin-deficient BMSC (**Fig. 5A-B**). Evidence emerges that high endoglin expression in stromal cells (BMSC or Cancer Associated Fibroblasts) supports glioma, breast and pancreatic cancer progression (47, 48) but the mechanisms at work are unknown. Here we established that high-stromal endoglin is required to support *in trans* the gain of tumor translational activity induced by stromal syntenin-deficiency (**Fig. 6A & 7C**). How syntenin deficiency leads to high cellular/membrane endoglin might be related to role of syntenin in late endosomal trafficking. Indeed, syntenin is well-known to address its PDZ interactors to the lumen of late endosomes for exosomal secretion (10, 16). Note that we here established that endoglin binds to the syntenin PDZ domains and that the two proteins colocalize intracellularly (see **Fig. 5D & 5E**). The increased cellular levels of endoglin observed upon syntenin loss-of-function might thus (at least in part) be due to defective targeting of endoglin to late endosomes/exosomal secretion. The much lower levels of endoglin accumulating into small EVs isolated from syntenin-deficient stromal cells (**Fig. 5C**) strongly supports this hypothesis. Secreted endoglin has recently been shown to exert anti-tumoral effect by working as decoy in TGFβ signaling (49). Whether a lack of vesicular endoglin secretion may contribute to the pro-tumoral effect of syntenin-deficient stroma remains to be clarified. Additionally or alternatively, as co-receptor for TGFβ, cell-associated endoglin will potentially support intracellular SMAD and AKT signaling (47). Noteworthy, the TGFβ receptor ligand BMP9 was recently shown to stimulate protein synthesis by inducing the activation of RPS6 (50). It would thus not be surprising that stromal endoglin and tumor TGFβ receptor cooperate *in trans* to potentiate AKT/RPS6 signaling and protein synthesis in AML cells.

Overall, our study underscores the importance of syntenin for cell-to-cell communication and uncovers unexpected molecular mechanisms that may govern tumor-stroma communication (**Fig. 7**).

## Materials and Methods

### Antibodies and reagents are listed in Supplementary Table S1

#### Cell lines

All human cell lines were obtained from the American Type culture collection (ATCC; Manassas, USA) and were grown in media supplemented with 10% heat-inactivated FBS (Eurobio, Les Ulis, France) and incubated at 37 °C, with 5% CO2. Human bone marrow stromal cell line HS5 was maintained in DMEM (Gibco, Carlsbad, USA). AML cell lines U937 (FLT3-negative, miR-155^low^) and HL60 (FLT3-WT, miR-155^low^)(24, 25) were maintained in minimum essential medium-α medium (Gibco, Carlsbad, USA). MOLM-14 (FLT3-ITD+, miR-155^high^)(51) were maintained in RPMI-1640 medium (Gibco, Carlsbad, USA). The cultured cells were split every 2–3 days, maintaining an exponential growth. FLB1 (miR-155^low^) primary mouse leukemia cells were provided by Pr. O.Hérault and were generated into C57BL/6J CD45.1 mice, as described previously (29).

#### Animal studies

CD45.2 C57BL/6J mice were purchased from Janvier Laboratories (France). CD45.2 C57BL/6J syntenin-knockout (synt-KO) animals were generated as previously described (52) and the hygromycin-resistance gene was removed using Flipper mice (53). Both male and female mice were used between 6 and 11 weeks of age and were housed under specific pathogen-free conditions. All experiments were performed in compliance with the laws and protocols approved by animal ethics committees (Agreement No. APAFIS#5123-2016041809182209v2). For *in vivo* expansion, freshly thawed 50,000 CD45.1 FLB1 cells were resuspended in PBS and injected into the retro-orbital vein of CD45.2 mice. Control mice consisted of PBS injected mice. AML development was monitored by flow cytometry, analyzing the percentage of CD45.1 positive cells in the blood. Recipient mice were sacrificed when blast levels reached >10% of total white blood cells in the peripheral blood (survival threshold), and leukemic cells were collected from femurs and tibias. For serial transplantation assays, blasts collected from the bone marrow (>90% blast invasion) of 3 different animals were pooled and used (50,000 cells/mouse) for re-injection in the following transplantation round. AML progression was assessed weekly, and animals were sacrificed at survival threshold.

#### Isolation and culture of murine bone marrow stromal cells

6 to 8 weeks old female CD45.2 C57BL/6J WT or synt-KO mice were sacrificed, hind limbs were collected and stripped of their skin and muscle manually. For isolation of the primary cells, the bone marrow from the femora and the tibia was flushed in RPMI buffer containing 1% FBS and 1% of penicillin-streptomycin (Gibco, Carlsbad, CA, USA). Red blood cells were removed using ACK lysis buffer (Gibco, Carlsbad, CA, USA) 10 min at room temperature. Cells were then washed twice into RPMI medium and seeded in a rat tail collagen-1 (5μg/cm^2^; Gibco, Carlsbad, CA, USA) coated flask for cell culture. Murine BMSC were routinely grown in RPMI medium supplemented with 10% FBS, 10% horse serum (Gibco, Carlsbad, CA, USA), 1%Penicillin-Streptomycin, and 1%Sodium-Pyruvate, in plastic flasks coated with collagen. The growing cells were characterized as murine bone marrow stromal cells (BMSC) by flow cytometry analysis: dead cells were excluded using DRAQ7 (Beckman Coulter, Brea, CA, USA) and cells were stained with antibodies for the positive (Sca1, CD51, CD106, CD105, CD54, CD44 & CD295) and the negative (CD45, CD3e, CD11c, B220, CD11b, CD19 & Ter119) selection of this lineage. Expanded BMSC WT/synt-KO were used at the fourth to the tenth passages.

#### Long-term co-culture experiments

Transfected HS5 and passage 4 to 10 murine BMSC (WT or synt-KO) were irradiated at 30 Gy prior co-culture to stop their proliferation during the co-culture assay. Irradiated murine BMSC (10,000) or HS5 cells (25,000) were seeded in 12-well plates and were cultured in RPMI medium containing exosome-depleted FCS (10%) for 24h. Then, AML cells (ratio 5:1 to stromal cells) were added and maintained in long-term co-culture for 1 month, with a media change every 3-4 days.

#### Western blots

The proteins were heat-denaturated in Laemmli sample buffer, fractionated in 12.5% or 15% gels by SDS–PAGE and electro-transferred to nitrocellulose membrane. Membranes were stained with Ponceau red and immunoblotted with the indicated primary antibodies (**Table S1**), and HRP-conjugated secondary antibodies (Mouse or Rabbit, Thermofisher scientific; 1/10000). Signals were visualized using Amersham ECL Prime Western Blotting Detection Reagent (GE Healthcare).

#### Mass Spectrometry Analysis

FLB1 CD45.1 cells were isolated and sorted by FACs after 4 rounds of serial transplantation from WT and synt-KO mice, using five animals for each condition. Two million of FLB1 CD45.1 cells per condition were lysed in RIPA buffer and analyzed by mass spectrometry as described in supplementary methods.

#### Measurement of protein synthesis

The alkyne analog of puromycin **“**O-propargyl-puromycin” (OPP) was purchased from Medchem source (Federal Way, WA, USA). For *in vivo* analysis, OPP (50 mg kg^−1^ body mass; pH 6.4–6.6 in PBS1X) was injected intraperitoneally. One hour later, mice were euthanized, and the BM was collected. To differentiate the CD45.1 FLB1 cells from the CD45.2 host cells, 3 × 10^6^ cells isolated from the BM were pre-stained with CD45.1 antibody at room temperature for 30min. Cells were then washed twice in Ca^2+^- and Mg^2+^-free PBS1X and fixed in 0.5 ml of 1% paraformaldehyde in PBS1X for 15 min on ice. Cells were washed in PBS1X, then permeabilized in 200 μl of PBS1X supplemented with 3% fetal bovine serum and 0.1% saponin (Sigma-aldrich, Saint-Louis, MO, USA) for 5 min at room temperature (20–25 °C). To visualize the incorporation of OPP into newly translated proteins, the azide-alkyne cycloaddition was performed using the Click-iT Cell Reaction Buffer Kit (Life Technologies, Carlsbad, CA, USA) and azide conjugated to Alexa Fluor 555 (Life Technologies, Carlsbad, CA, USA) at 5 μM final concentration. After the 30-min reaction, the cells were washed twice in PBS1X supplemented with 3% of FBS and 0.1% saponin, then resuspended in PBS1X supplemented with 4′,6-diamidino-2-phenylindole (DAPI; 4 μg ml^−1^ final concentration) and analyzed by flow cytometry. Relative rates of protein synthesis were calculated by normalizing OPP signals to whole bone marrow after subtracting autofluorescence background. For *in vitro* analysis, 0.5 × 10^6^ cells were seeded in 2ml of RPMI medium containing FCS (10%). OPP (10 μM final concentration) was added to the culture medium for 1 h. Control cells consisted of PBS1X treated cells. The cells were collected, fixed, permeabilized, and the azide-alkyne cycloaddition was performed as described above. Mean OP-Puro fluorescence reflected absolute fluorescence values for each cell population from multiple independent experiments.

#### Statistical analysis

Differences between groups were determined by using 1-way or 2-way analysis of variance (ANOVA), followed by a Bonferroni post-test using GraphPad Prism v8.0c software. Single comparisons were carried out using the parametric Student t-test or the nonparametric Mann-Whitney U test. P < 0.05 was considered statistically significant.

## Supporting information

Supplementary informations

## Acknowledgments

We thank Prof. Olivier Herault (University of Tours, France) for providing us with FLB1 cells, Prof. Peter Carmeliet (University of Leuven, Belgium) for providing us with FLPeR mice, Profs. Ingrid Struman and Pierre Close (GIGA, University of Liège, Belgium) for helpful exchanges about miRNA biology and protein synthesis, respectively, and Dr. Jean-Hugues Guervilly for its scientific advice. We are grateful to the core flow cytometry and the animal facility of the CRCM for providing services. This work was supported by the Concerted Actions Program of the KU Leuven (GOA/12/016), the internal funds of the KU Leuven (C14/20/105), the Belgian Foundation against cancer (STK, FA/2014/294), the French Foundation for Cancer Research (ARC, PJA 20141201624), the Institut National du Cancer (INCa, subvention 2013-105), the National Research Agency (ANR, Investissements d’Avenir, A*MIDEX project ANR-11-IDEX-0001-02), the Institut National de la Santé et de la Recherche Médicale (INSERM), and by the Fund for Scientific Research—Flanders (FWO, G.0846.15 and G0C5718N). This study was partly supported by research funding from the Canceropôle Provence Alpes Côte d’Azur, Institute for Cancer and Immunology (Aix-Marseille University), Institut National du Cancer and Région Sud. RL was the recipient of fellowships from the ARC French Foundation for Cancer Research.

