## Supplementary Figures for "Downregulation of stromal syntenin sustains AML development"

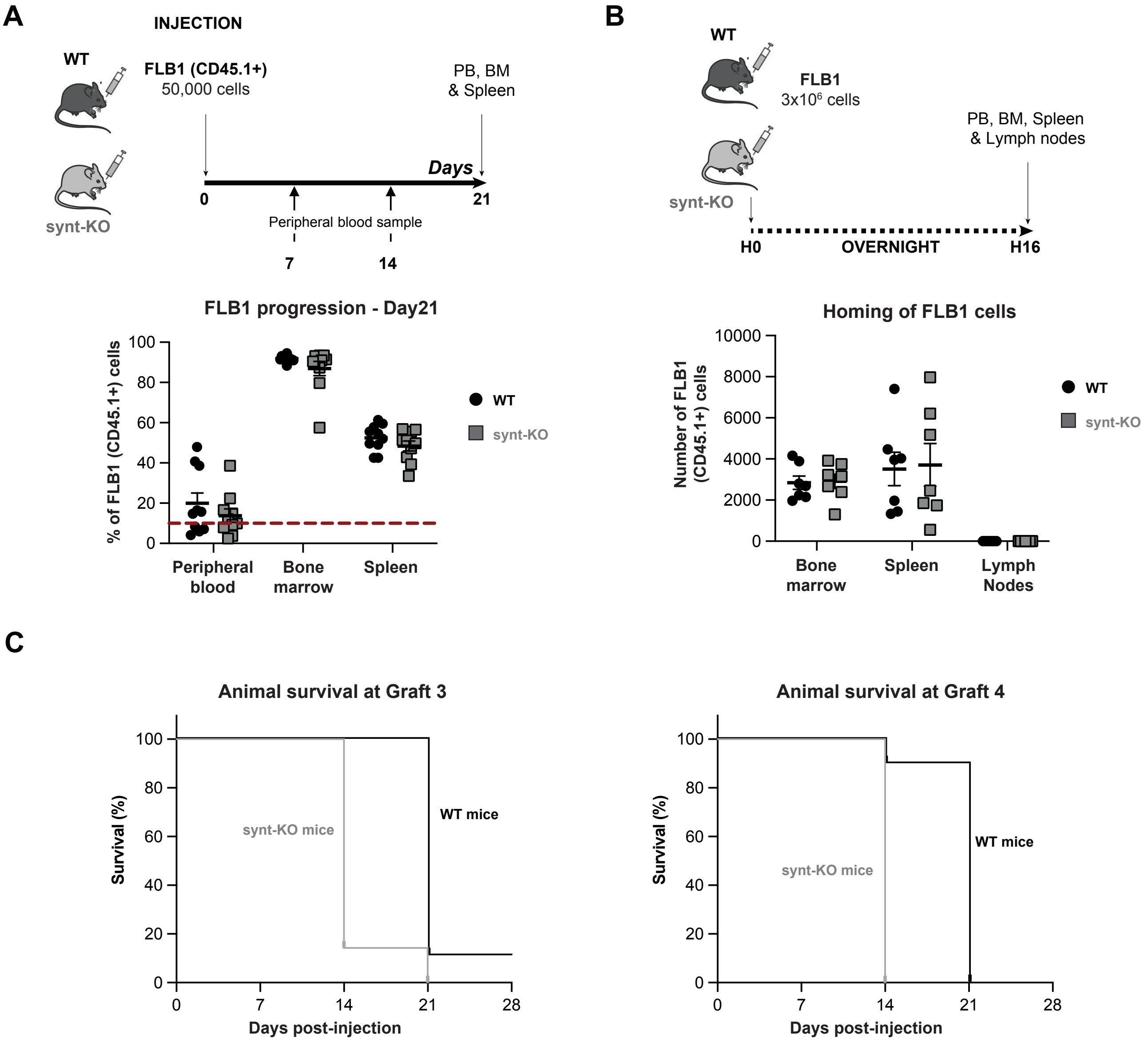
A
INJECTION
WT
FLB1 (CD45.1+)
50,000 cells
PB, BM
& Spleen
Days
0
21
synt-KO
Peripheral blood sample
7
14
FLB1 progression - Day21
% of FLB1 (CD45.1+) cells
100
80
60
40
20
0
Peripheral blood
Bone marrow
Spleen
WT
synt-KO
B
WT
FLB1
3x10[6] cells
PB, BM, Spleen
& Lymph nodes
synt-KO
H0
OVERNIGHT
H16
Homing of FLB1 cells
Number of FLB1 (CD45.1+) cells
10000
8000
6000
4000
2000
0
Bone marrow
Spleen
Lymph Nodes
WT
synt-KO
C
Animal survival at Graft 3
Survival (%)
100
80
60
40
20
0
synt-KO mice
WT mice
0
7
14
21
28
Days post-injection
Animal survival at Graft 4
Survival (%)
100
80
60
40
20
0
synt-KO mice
WT mice
0
7
14
21
28
Days post-injection

**Figure S1**

**A**

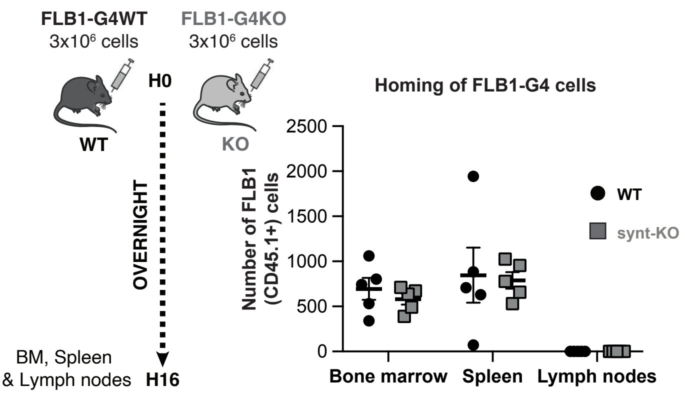


**B**

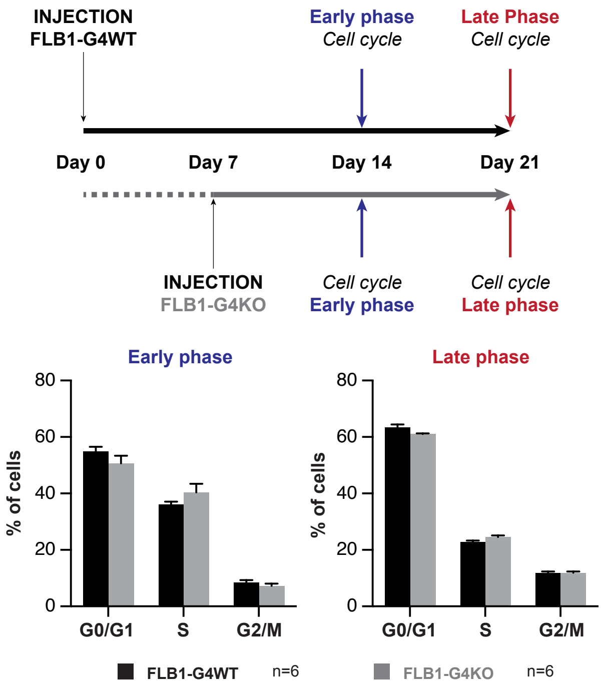


**C**

| Group | Dose (cells) | Mice tested | Mice engrafted (% of FLB1>1% in PB) | LIC frequency & confidence interval [lower - upper] |
|---|---|---|---|---|
| FLB1-G4WT | 2000 | 6 | 5 | 1/401 [691 - 233] |
| | 1000 | 6 | 5 | |
| | 500 | 6 | 5 | |
| | 200 | 12 | 8 | |
| | 20 | 6 | 1 | |
| FLB1-G4KO | 2000 | 6 | 6 | 1/365 [630 - 212] |
| | 1000 | 6 | 6 | |
| | 500 | 6 | 4 | |
| | 200 | 12 | 5 | |
| | 20 | 6 | 0 | |

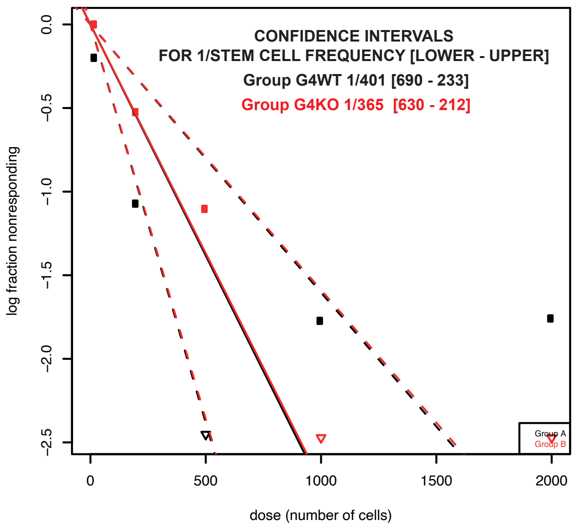


**D**

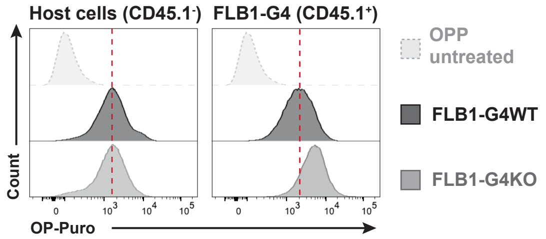


**E**

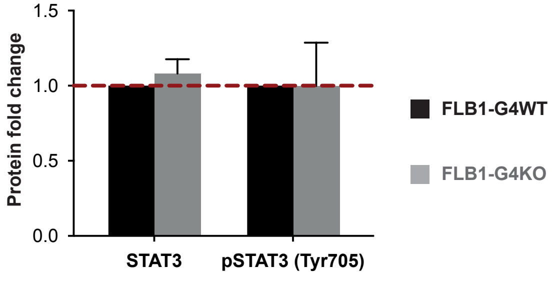


**Figure S2**

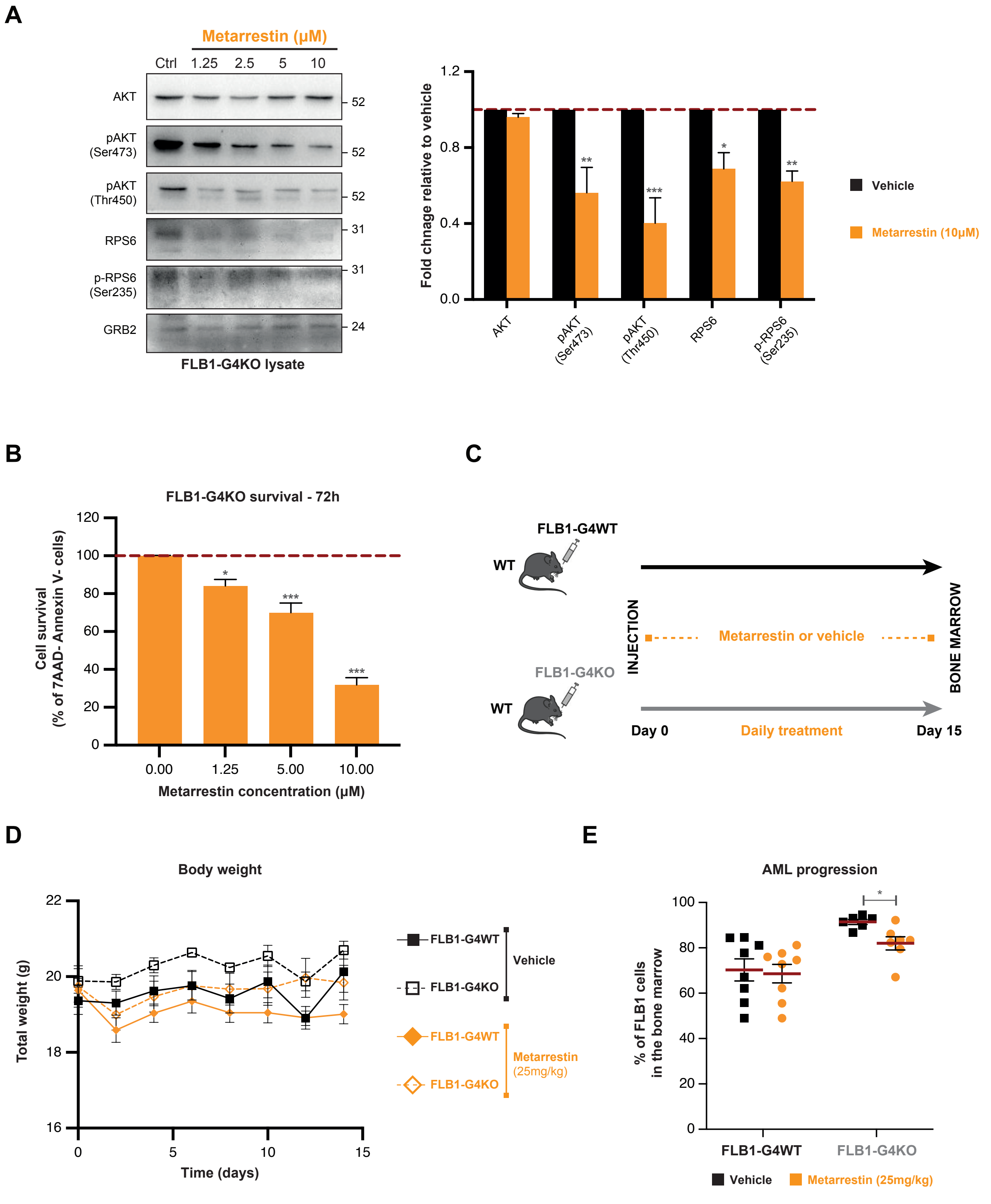
A
Metarrestin (μM)
Ctrl 1.25 2.5 5 10
AKT
pAKT (Ser473)
pAKT (Thr450)
RPS6
p-RPS6 (Ser235)
GRB2
52
52
52
31
31
24
FLB1-G4KO lysate
Fold chnage relative to vehicle
1.2
0.8
0.4
0.0
AKT
pAKT (Ser473)
pAKT (Thr450)
RPS6
p-RPS6 (Ser235)
Vehicle
Metarrestin (10μM)
B
FLB1-G4KO survival - 72h
Cell survival (% of 7AAD- Annexin V- cells)
120
100
80
60
40
20
0
0.00
1.25
5.00
10.00
Metarrestin concentration (μM)
C
FLB1-G4WT
WT
FLB1-G4KO
WT
INJECTION
Metarrestin or vehicle
BONE MARROW
Day 0
Daily treatment
Day 15
D
Body weight
Total weight (g)
22
20
18
16
0
5
10
15
Time (days)
FLB1-G4WT
FLB1-G4KO
Vehicle
FLB1-G4WT
FLB1-G4KO
Metarrestin (25mg/kg)
E
AML progression
% of FLB1 cells in the bone marrow
100
80
60
40
20
0
FLB1-G4WT
FLB1-G4KO
Vehicle
Metarrestin (25mg/kg)

Figure S3

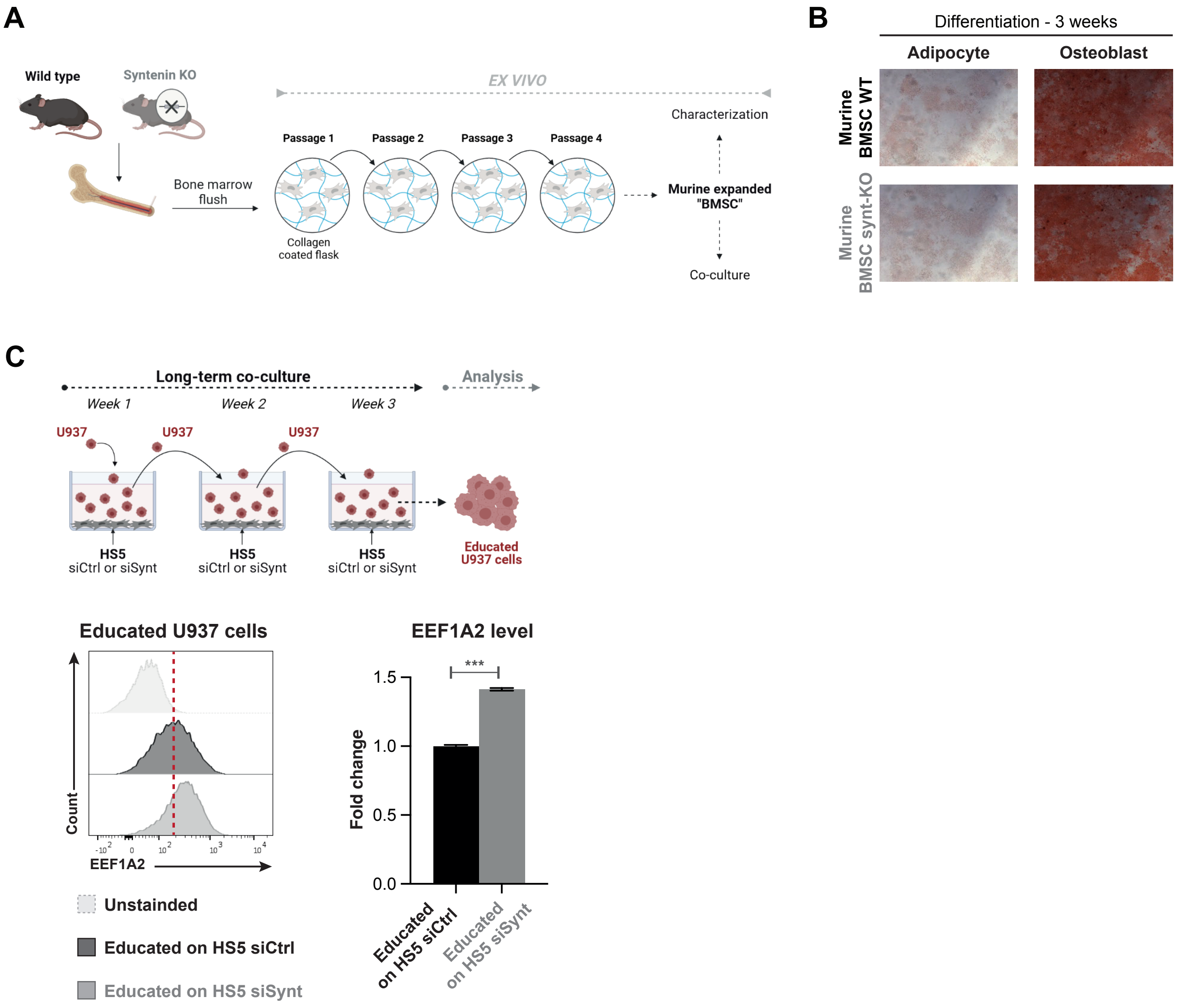
A
Wild type
Syntenin KO
EX VIVO
Characterization
Passage 1
Passage 2
Passage 3
Passage 4
Bone marrow flush
Collagen coated flask
Murine expanded "BMSC"
Co-culture
B
Differentiation - 3 weeks
Adipocyte
Osteoblast
Murine BMSC WT
Murine BMSC synt-KO
C
Long-term co-culture
Analysis
Week 1
Week 2
Week 3
U937
U937
U937
HS5 siCtrl or siSynt
HS5 siCtrl or siSynt
HS5 siCtrl or siSynt
Educated U937 cells
Educated U937 cells
Count
EEF1A2
Unstainded
Educated on HS5 siCtrl
Educated on HS5 siSynt
EEF1A2 level
***
Fold change
1.5
1.0
0.5
0.0
Educated on HS5 siCtrl
Educated on HS5 siSynt

**Figure S4**

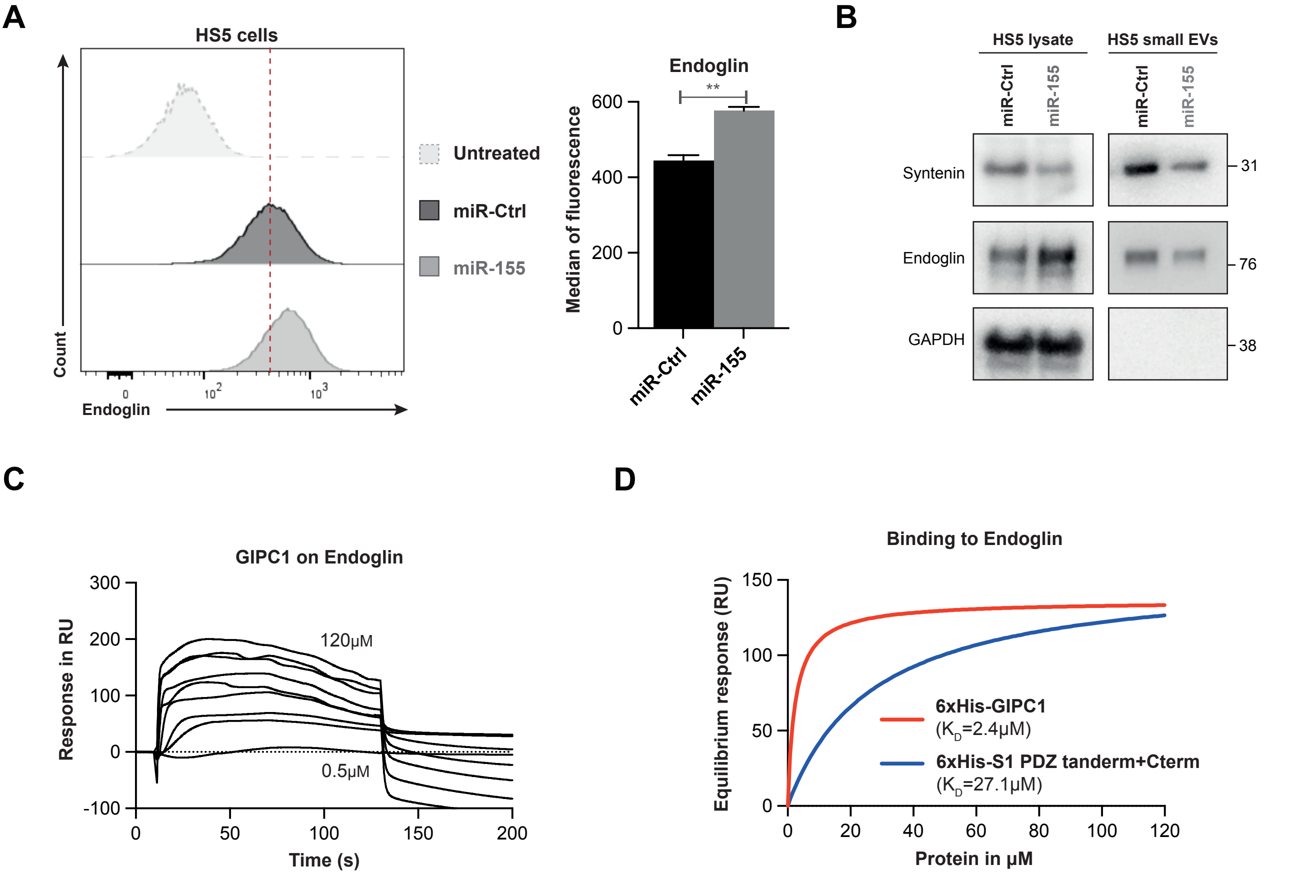

A
HS5 cells
Count
Endoglin
0
10^2
10^3
Untreated
miR-Ctrl
miR-155
Endoglin
Median of fluorescence
600
400
200
0
**
miR-Ctrl
miR-155
B
HS5 lysate
HS5 small EVs
miR-Ctrl
miR-155
miR-Ctrl
miR-155
Syntenin
Endoglin
GAPDH
31
76
38
C
GIPC1 on Endoglin
Response in RU
300
200
100
0
-100
120μM
0.5μM
0
50
100
150
200
Time (s)
D
Binding to Endoglin
Equilibrium response (RU)
150
100
50
0
6xHis-GIPC1
(KD=2.4μM)
6xHis-S1 PDZ tanderm+Cterm
(KD=27.1μM)
0
20
40
60
80
100
120
Protein in μM


**Figure S5**

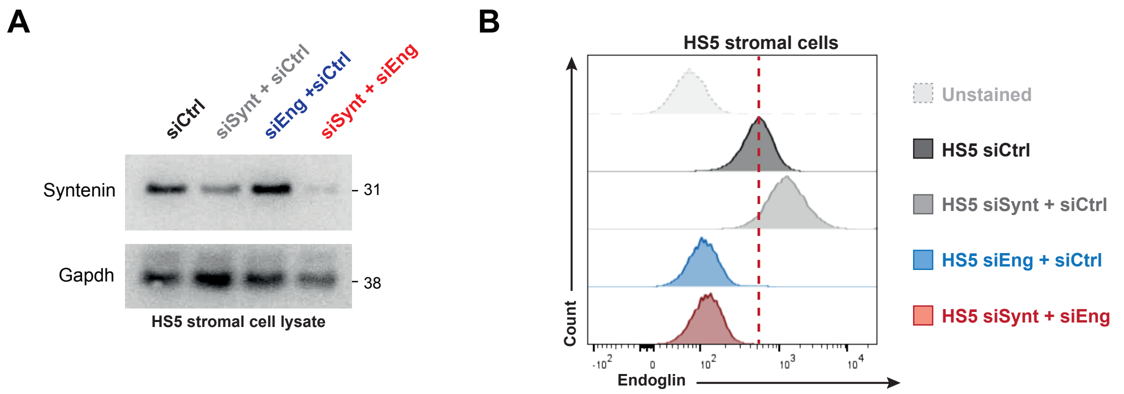
A
siCtrl
siSynt + siCtrl
siEng +siCtrl
siSynt + siEng
Syntenin
31
Gapdh
38
HS5 stromal cell lysate
B
HS5 stromal cells
Count
Endoglin
Unstained
HS5 siCtrl
HS5 siSynt + siCtrl
HS5 siEng + siCtrl
HS5 siSynt + siEng

**Figure S6**
