## Supplementary informations for "Downregulation of stromal syntenin sustains AML development"

**Supplementary Materials and Methods**

**Gene expression data analysis.** To assess the expression of syntenin in BMSC from mice inoculated with different subtypes of leukemia, we collected gene expression profiling data from the National center for Biotechnology Information (NCBI, https://www.ncbi.nlm.nih.gov) with the accession number GSE97194. As previously described (1), murine AML cells carrying different gene rearrangements (MLL-ENL, AML1-ETO9a or MLL-ENL + FLT3-ITD) were transplanted into C57Bl/6 mice (6-8weeks old). PBS was injected in the control group. Transplanted leukemia was allowed to progress for 3-4 weeks according to the disease burden. At last, mice were sacrificed and BMSC were isolated by FACS using a combination of cell surface markers (Ter119$^-$, CD31$^-$, CD45$^-$, Sca1$^+$, CD106$^+$, CD105$^+$ and CD140$^+$). RNA was extracted, amplified and biotinylated as previously described (1). Microarray data were generated using IlluminaHT12 version 4 mouse whole genome arrays. Data extraction and analysis was performed using the webserver GEOexplorer (https://geoexplorer.rosalind.kcl.ac.uk), as previously described (2). Briefly, normal BMSC (Ctrl_expt, n=3) were compared to AML-BMSC (MLL&FLT3-ITD_p53WT_Expt1, MLL&FLT3-ITD_p53-/-_Expt1; AML-ETO_Expt, n=2; MLL-ENL_Expt; n=2) for differential gene expression analysis. Data were normalized and p-values were adjusted using Benjamini & Hochberg procedure (**Supplementary Table S2 & S3**). The gene that are differentially expressed were selected based on a p-value P<0.01 and a difference >2.

**Cell transfection.** For RNAi experiments, cells at a confluence of 50% were transfected with 30nM RNAi using Lipofectamine RNAiMAX reagent (Life technologies, USA); cells were analyzed 48 to 96 hours after RNAi treatment. RNAi targeting human SDCBP (5'-GCAAGACCUUCCAGUAUAA-3'), a SMART pool targeting human ENG and the non-targeting control RNAi (siCNT) were purchased from Dharmacon Inc. For miRNA treatment, HS5 cells were transiently transfected using 30nM of hsa-miR-155-5p mimic or 50nM of mirVana negative control, in presence or absence of 30nM of antagomiR-155 (miR-155 inhibitor). Expanded murine BMSC WT were transiently transfected using 30nM of mmu-miR-155-5p mimic or 50nM of mirVana negative control.

**Mass Spectrometry Analysis and Protein Quantification.**

***- Sample preparation and labelling.*** Equal amounts of protein (i.e. around 100µg) were aliquoted and adjusted to 100 µl using 100 mM TEAB. Proteins were reduced using TCEP and then alkylated using iodoacetamide as mentioned in the TMT mass tag labelling instructions (Thermo Scientific). Proteins were precipitated using six volumes of cold acetone at -20°C overnight. The pellet was then suspended in 100 mM TEAB and digested with 2.5µg of Trypsin (Promega), overnight at 37°C. Peptides were labelled with TMT reagents as described in the TMT labelling instructions (Thermo Scientific). Briefly, samples were transferred to TMT reagents reconstituted using 40ul of anhydrous acetonitrile and then incubated at room temperature for one hour. Five biological replicates for both conditions, i.e. PWT and PKO FLB1, were distributed in the ten TMT channels. The reaction was quenched by adding 8 µl of 5% hydroxylamine and incubating for 15 min. Labelled peptides were mixed, and dried to remove organic solvent prior to clean-up via Sep-Pak (50 mg C18 SepPak; Waters). Labeled peptide mixtures were separated in 80 fractions via high-pH reversed phase chromatography (Waters; Xbridge C18 analytical column) and concatenated into thirteen fractions. Samples were dried and stored at −80 °C prior to LC-MS analysis.

***- Mass spectrometry analysis.*** Samples were reconstituted in 4% acetonitrile/0.1% trifluoroacetic acid prior to analysis by liquid chromatography (LC)-tandem mass spectrometry (MS/MS), using an Orbitrap Fusion Lumos Tribrid Mass Spectrometer (Thermo Fischer) online with an Ultimate 3000RSLCnano chromatography system (Dionex). Peptides were separated on a reverse phase LC EASY-Spray C18 column from Dionex (PepMap RSLC C18, 50 cm x 75 µm I.D, 100 Å pore size, 2 µm particle size) at 300 nL/min flow rate and 40°C and using a two steps linear gradient (4-20% acetonitrile/H20; 0.1 % formic acid for 180 min and 20-45-45% acetonitrile/H2O; 0.1 % formic acid for 60 min. An EASY-Spray nanosource was used for peptide ionization (2 200 V, 275°C). The mass spectrometer was used in data dependent mode to switch consistently between MS1, MS2 for peptide identification and multinotch-MS3 for protein abundance measurements. Time between Masters Scans was set to 3 seconds. MS1 spectra were acquired with the Orbitrap in the range of m/z 375-1500 at a FWHM resolution of 120 000 measured at 200 m/z. AGC target was set at 4.0x10E5 with a 50 ms maximum injection time. The more abundant precursor ions were selected for MS2 (Top speed 3 seconds) and collision induced dissociation fragmentation at 35% was performed and analysed in the linear ion trap using the "Inject Ions for All Available Parallelizable time" option with a maximum injection time of 105 ms and an AGC target of 1.0x10E5. For Multi-Notch MS3, the top 10 precursor ions from each MS2 scan were fragmented by HCD followed by orbitrap analysis (Resolution = 60 000; First mass = 100 m/z; AGQ target = 250000; MaxIT = 86ms, 100-500 m/z). Charge state screening was enabled to include precursors with 2 and 7 charge states. Dynamic exclusion was enabled with a repeat count of 1 and a duration of 60s. The mass spectrometry proteomics data have been deposited to the ProteomeXchange Consortium via the PRIDE partner repository with the data set identifier PXD023602.

***- Database search and relative quantification.*** Relative TMT intensity-based quantification was processed using the freely available MaxQuant computational proteomics platform, version 1.6.3.4. Raw data were searched against the mouse database, extracted from UniProt on the 9th of December 2019 and containing 20379 entries (reviewed) supplemented with known contaminants. Searches were performed with default parameters trypsin enzymatic cleavage with up to two missed cleavages, and three variable modifications allowed per peptide. Searches were performed with variable methionine oxidation (+15.99491), static cysteine carboxyamidomethylation (+57.02146), and static TMT modifications on lysine and the peptide N-termini (+229.16293). The false discovery rate (FDR) at the peptide and protein levels were set to 1% and determined by searching a reverse database. For protein grouping, all proteins that cannot be distinguished based on their identified peptides were assembled into a single entry according to the MaxQuant rules.

TMT reporter intensities were corrected for isotope impurities according to the lot used. The statistical analysis was done with Perseus program (version 1.6.2.1). First corrected TMT reporter intensities were base 2 logarithmized to obtain a normal distribution and normalized by subtracting the median for each TMT channel. The few proteins with missing values were removed from the dataset. To determine whether a given detected protein was specifically differential, a two-sample t-test was done using permutation-based FDR-controlled at 0.01 and employing 250 permutations. The p value was adjusted using a scaling factor s0 with a value of 0.4 for the FLB1-G4WT/-G4KO with n=5.

***In vitro* differentiation of murine expanded BMSC*.*** Osteoblastic differentiation was induced by culturing expanded BMSC, WT/synt-KO, for 4 weeks with 50 µg ml−1 of l-ascorbic acid 2-phosphate, 10 mM glycerophosphate (Sigma-aldrich, Saint-Louis, MO, USA) and 15% FBS in α-MEM supplemented with penicillin-streptomycin. Adipocyte differentiation was induced with 1 µM dexamethasone, 10 µg ml−1 of insulin (Sigma-Aldrich, Saint-Louis, MO, USA) and 10% FBS in α-MEM with penicillin-streptomycin. All cultures were maintained with 5% CO2 in a water-jacketed incubator at 37 °C, and media were replaced every 2-3 days. To assess *in vitro* differentiation into mesenchymal lineages, cells were briefly washed with phosphate buffered saline solution (PBS1X; Gibco, Carlsbad, CA, USA) and fixed for 10 min at room temperature with 4% paraformaldehyde (Santa Cruz). For the detection of mineral calcium deposition, osteoblasts were stained with Alizarin Red solution (Sigma-Aldrich, Saint-Louis, MO, USA) for 30min and rinsed 3 times with distillated water. Adipocytes were stained with Oil Red O (Sigma-Aldrich, Saint-Louis, MO, USA) as follows: cells were washed with 60% isopropanol and allowed to dry completely. Oil Red O working solution was prepared as a 6:4 dilution in distilled water of a 0.35 g ml−1 Oil Red O solution in isopropanol (Sigma-aldrich, Saint-Louis, MO, USA) and filtered 20 min later. Cells were incubated for 60 min with Oil red O working solution and rinsed four times using distilled water.

***In vivo* homing assays.** For homing assays, a total of $3\times10^6$ FLB1 (CD45.1) cells was injected into the retro-orbital vein of WT or synt-KO CD45.2 C57Bl/6J mice. 16 hours after cell inoculation, animals were sacrificed, bone marrow, spleen and lymph nodes were collected for analysis. Numbers of FLB1 in the bone marrow and the spleen were evaluated by flow cytometry using CD45.1/CD45.2 antibodies and CountBright™ Absolute Counting Beads according to the manufacturer.

***In vivo* limiting dilution.** Extreme Limiting Dilution Analysis (ELDA) comparing populations for enrichment in stem cell was performed as previously described (PMID: 19567251). Five doses (20, 200, 500, 1000, 2000) of CD45.1 FLB1-G4 cells, isolated from WT or synt-KO animals at graft 4, were transplanted into CD45.2 C57Bl/6J mice. The cut-off for engraftment was set at FLB1 (CD45.1) cells composing more than 1% of the total white blood cells in the peripheral blood, up to 12 weeks after cell inoculation. Leukemia initiating cell (LIC) frequencies were determined using the webtool interface "http://bioinf.wehi.edu.au/software/elda/".

**Flow Cytometry.** For extracellular staining, cells were incubated with appropriate antibodies (**Table S1**) diluted in PBS1X supplemented with 0.1% of FBS to block non-specific antibody binding, for 30 min at room temperature. Intracellular staining for indicated markers was performed using BD Cytofix/Cytoperm™ Fixation/Permeabilization kit (BD Biosciences; **Table S1**). For absolute counting of cells, 10µL of CountBright absolute counting beads (ThermoFisher Scientific) were added to the stained cells just before acquisition on the flow cytometer. The acquisitions were performed by a BD LSRII flow cytometer or a BD Fortessa flow cytometer (BD Biosciences). Data analysis was conducted with FlowJo software (BD).

**AML cell viability.** AML blasts were collected, and early/late apoptosis was measured by using the PE Annexin V apoptosis detection kit with 7-AAD (Biolegend, San Diego, USA) according to the manufacturer. Annexin V positive but 7-AAD negative (early apoptotic cells) and both Annexin V and 7-AAD positive cells (late-stage apoptosis) were determined by using FACS LSRII flow cytometer and data were analyzed with Flowjo software (BD Biosciences, Franklin Lakes, USA).

**Exosomes and total cell lysates.** For comparative analyses, exosome-enriched fractions were collected from equivalent amounts of culture medium, conditioned by equivalent amounts of HS5 cells transfected with siRNA targeting syntenin (siSynt) or control siRNA (siCtrl). After 16hours, cell media were collected and small extracellular vesicles were isolated by three sequential centrifugation steps at 4 °C: 10 min at 500×g, to remove cells and large debris; 30 min at 10,000×g, to remove large extracellular vesicles; and 1h30min at 100,000×g, to pellet small extracellular vesicles (exosome-enriched fraction), followed by one wash with 1400µL of PBS1X (100,000×g, 1h), to remove soluble serum and secreted proteins. Small extracellular vesicle pellets were then re-suspended in 100µL of PBS1X. The lysates from corresponding cultures were cleared by centrifugation at 1500rpm for 5 min and then resuspended in lysis buffer (TrisHCL pH 7.4 30mM, NaCl 150mM, 1% NP40 (IGEPAL), 1µg/ml aprotinin, 1µg/ml leupeptin). Although only small variations were observed from sample to sample, exosomal amounts loaded for the western blot were normalized according to the number of parent cells from where exosomes were secreted.

**Reverse transcription-polymerase chain reaction (RT-PCR).** RNA was extracted from cells using Nucleospin RNA isolation kit (Macherey-Nagel). Reverse Transcription was done using PrimeScript RT Reagent with gDNA Eraser kit (TAKARA CLONTECH) and MJ Research PTC-220 Dyad PCR System (Conquer Scientific). Quantitative PCR reactions were performed using SsoAdvanced Universal Sybr Green Supermix kit (BIORAD) and analyzed using CFX96 Touch Real-Time PCR Detection System (BIORAD). The expression level was normalized to the housekeeping L32 and GAPDH genes. The cDNAs were amplified by 35 cycles with sets of specific primers (**Supplementary Table S4**). Each PCR cycle consisted of 15s of denaturation at 95°C, 30s of annealing and extension at 60°C. Quantifications were normalized to corresponding L32 and GAPDH RNA values and expressed as relative expression using the $2(-\Delta\Delta CT)$ method (3).

**Cell cycle assays.** Distribution of cell cycle phases was determined by Propidium Iodide (PI; Life Technologies, Carlsbad, CA, USA) staining and flow cytometry analysis. Briefly, $5\times10^6$ FLB1-G4 (CD45.1) cells were isolated from the bone marrow of WT or Synt-KO mice and stained with CD45.1 antibody. Cells were fixed in 1 mL ice-cold 70% ethanol for 30min at 4°C and stored at −20 °C. At the time of analysis, the cells were centrifugated, washed once again in PBS1X and stained with a freshly made solution containing 40μg/ml of PI, 0.1% Triton x-100 and 40μg/ml of RNaseA (Life Technologies, Carlsbad, CA, USA) in PBS1X. Percentages of cells within cell cycle compartments (G1, S and G2/M) were determined using a FACS LSRII flow cytometer and data were analyzed with Flowjo software.

**Surface plasmon resonance experiments.** SPR measurements were carried out at 25°C using a BIAcore T200 instrument (GE Healthcare). A total of 100 resonance units (RU) of biotinylated ligands corresponding to the last 25 amino acids of endoglin wild-type (SSESSSTNHSIGSTQSTPCSTSSMA) were immobilized on a streptavidin-sensor chip (GE Healthcare). A 6xHis-tagged syntenin construct (containing the two PDZ domains and the C-terminal domain) or 6xHis-tagged GIPC1 construct were perfused at 30 µl per min in running buffer (10mM HEPES pH 7.4, 150mM NaCl and 0.005% Tween 20) at different concentrations (µM). Recombinant proteins were prepared as described previously (PMID: 27386966). The reference channel was blank immobilized. The injection time was 120s, allowing binding to reach equilibrium. The dissociation time was 90s. The surfaces were regenerated between runs, by short pulses of 50mM NaOH and 1M NaCl at 30 µl per min flow rate. Sensorgrams were corrected for binding to reference surfaces and for buffer effects (blank subtracted) before further data analysis. Apparent equilibrium dissociation constants ($K_D$) were calculated by fitting the data to a simple Langmuir binding isotherm using GraphPad Prism. RUs at equilibrium were plotted as a function of protein concentration.

**Immunofluorescence staining and confocal microscopy.** HS5 cells were treated with siRNA targeting syntenin (siSyntenin) or control (siCNT) for 48h. After 48h, cells were fixed with 100% Methanol for 5 min, washed in PBS and permeabilized with 0.1% Triton X-100 for 10 min. Cells were then incubated with the indicated antibodies in PBS buffer containing 1% BSA. Cells were mounted in ProLong™ Diamond Antifade Mountant containing DAPI and observed with a Zeiss confocal microscope (LSM 880, Zeiss, France) with the corresponding lasers and 40× objectives. Confocal images were analyzed using Photoshop (Adobe, San Jose, CA) software.

**Supplementary figures**

**Fig. S1 (related to figure 2). Invasion and colonization of hematopoietic tissues (peripheral blood, bone marrow, spleen and lymph nodes) by FLB1 upon first transplantation (A,B) and animal survival upon serial transplantation (C), in wild-type versus syntenin-deficient animals. (A)** Upper part, scheme for the invasion assay. Non-irradiated 8-11w old wild-type (WT) and syntenin-knock-out (synt-KO) mice were injected with 50,000 FLB1 cells in the retro-orbital vein. Lower part, FACS analysis monitoring leukemia progression at day 21 in different tissues (as indicated). The dot-plot summarizes the percentage of CD45.1 FLB1 cells, relative to the total number of CD45+ cells ± SEM, in the different organs. **(B)** Upper part, scheme for the homing assay. Non-irradiated 8-10wks old WT and KO mice were injected with $3x10^6$ FLB1 cells in the retro-orbital vein. 12 hours after cell injection, animals were sacrificed and BM, spleen and lymph nodes were collected for analysis. Lower part, FACS analysis monitoring the number of FLB1 cells (using FITC-conjugated CD45.1-antibody) in different WT and synt-KO tissues as indicated. The dot-plot indicates mean values ± SEM. **(C)** Kaplan-Meier-like plot related to **fig. 1B**, comparing the survival of WT and synt-KO mice at graft 3 and 4. All animals were sacrificed when blast levels reached >10% in the peripheral blood. The survival curves for WT and synt-KO animals at graft 3 and 4 are significantly different using the log-rank test (***; P<0.0001).

**Fig. S2. (related to figure 3). FLB1-G4KO display similar homing, cell cycling and clonogenic properties as FLB1-G4WT. (A)** Left, scheme of the homing assay. Non-irradiated 8-10wks old WT and Synt-KO mice were injected with $3x10^6$ FLB1-G4WT or FLB1-G4KO cells in the retro-orbital vein. 16 hours after cell injection, animals were sacrificed and BM, spleen and lymph nodes were collected for analysis. Numbers of FLB1 cells in the BM and the spleen were counted by FACS, using the CD45.1-FITC antibody and CountBright™ Absolute Counting Beads according to the manufacturer. Results are expressed as mean number of CD45.1 cells per organ ± SEM. No signal was observed in lymph nodes. **(B)** Extreme Limiting Dilution Assay (ELDA) was performed to compare the frequency of Leukemia Initiating Cells (LIC) in FLB1-G4WT and FLB1-G4KO. Upper part, table illustrating the ELDA performed by injecting decreasing numbers of sorted FLB1-G4WT or FLB1-G4KO cells, as indicated in the "dose column". The mice were considered as engrafted when FLB1 (CD45.1) cells were detectable in the peripheral blood and composed at least 1% of the total cells in that compartment, with the numbers of successful engraftments indicated in the column "mice engrafted". LIC frequencies for FLB1-G4WT and FLB1-G4KO were estimated, as well as the interval of confidence for each condition. Lower part, log-fraction plot illustrating the ELDA fitted to FLB1-G4WT vs FLB1-G4KO. The slope of the line represents the estimated LIC frequency. LIC frequencies of FLB1-G4WT (black line) and FLB1-G4KO (red line) are respectively

1/401 and 1/365. The dotted (black and red) lines indicate the 95% confidence intervals. **(C)** Upper part, scheme for the cell-cycling assay. FLB1-G4WT cells were injected at day 0 into WT mice, while the injection of FLB1-G4KO cells into synt-KO mice (evolving more rapidly) occurred one week later. Cell cycling was analyzed at both early (day 14) and late (day 21) phases of leukemia progression. Animals were sacrificed and their BM was stained with propidium iodide. Lower part, the cell cycle stage of the CD45.1+ cells was then analyzed by FACS. Results are expressed as percentage of cells in the different phases of the cell cycle ± SEM. **(D)** Representative histograms showing OPP incorporation into host cells ($CD45.1^-$, left) and FLB1-G4 cells ($CD45.1^+$, right) isolated from the BM of WT (black histogram), synt-KO (dark gray histogram) mice, or from the BM of a PBS-injected control mouse (OPP untreated; light gray histogram). Note the higher incorporation of OPP in FLB1-G4KO compared to FLB1-G4WT. **(E)** Histogram representing mean signal intensities ± SEM, relative to signals obtained with FLB1-G4WT lysates, calculated from the analysis of 3 independent mice. Note, the absence of significant difference. Statistical analysis was performed using the nonparametric Mann-Whitney U test.

**Fig. S3 (related to figure 3). The EEF1A2 inhibitor Metarrestin diminishes the *in vitro* survival and in vivo aggressiveness of leukemia blasts educated by a syntenin-deficient host. (A)** Left, Western blots of total cell lysates from FLB1-G4KO treated with DMSO (Ctrl) or metarrestin (1.25 to 10µM) for 16h. Different markers of cell survival were analyzed, as indicated. GRB2 was used as control. Right, histograms representing the mean signal intensities in drug-treated cells relative to the signals in FLB1-G4KO cells treated with vehicle (DMSO) ± SEM, calculated from the analysis of 3 independent experiments. Statistical analysis was performed using the two-way analysis of variance (ANOVA) (*, P < 0.05; **P < 0.005; ***P < 0.0005). **(B)** Effect of Metarrestin treatment on FLB1-G4KO cell survival as determined in FACS. Cell survival in the presence of increasing concentrations of metarrestin was evaluated at 72h. Data represent the mean percentages of living FLB1-G4KO (CD45.1) cells ± SEM calculated from 3 independent experiments performed in triplicate. Statistical analysis was performed using the two-way analysis of variance (ANOVA) (*P < 0.05; ***P < 0.0005). **(C)** Scheme of the in vivo Metarrestin experiments. FLB1-G4WT cells were injected at day 0 into WT mice, while the inoculation of FLB1-G4KO cells into KO mice was delayed for one week, in order to synchronize the treatment. Animals were daily treated with Metarrestin or vehicle. Leukemia progression was assessed weekly, by FACS analysis as previously reported. **(D)** Daily evaluation of the body weight of each animal, treated with Metarrestin or vehicle. **(E)** FLB1 cell (CD45.1) frequencies in bone marrow aspirates were measured 15 days after cell inoculation. Results are expressed as mean percentage of FLB1 cells in the bone marrow ± SEM. Statistical analysis was performed using 1way ANOVA test (*, P < 0.05).

**Fig. S4 (related to figure 4). Murine BMSC isolation and characterization, and co-culture**

**experiments of HS5 human stromal cells with the U937 human leukemia model. (A)** Scheme illustrating the protocol for the preparation of murine expanded BMSC. 5-6w old WT and synt-KO mice were sacrificed, hind limbs were collected, and bones were flushed. Whole BM was plated on collagen coated flasks and expanded *ex vivo*. Hematopoietic cells were gradually eliminated by several washes, medium changes and at least four passages. **(B)** Characterization of the differentiation potential of BMSC. Expanded BMSC, WT and synt-KO, at passage 4 were cultivated under specific osteogenic or adipogenic conditions for 21 days. Alizarin Red and Red oil staining were performed to assess, respectively, Ca2++ deposition after osteogenic induction and lipid droplet accumulation after adipocyte differentiation. Note the similar differentiation profiles of WT and synt-KO BMSCs. **(C)** Upper, scheme for co-culture experiments using human leukemia and stroma model cells. HS5 (stroma) cells were transfected with siRNA targeting syntenin (siSynt) or control (siCtrl). After 24h, U937 (leukemia) cells were put in contact with HS5 cells. After one week of co-culture, U937 cells were collected and re-seeded on freshly transfected HS5 cells. After 3 weeks, U937 cells 'educated' by HS5-siCtrl or HS5-siSynt were collected for FACS analysis (bottom panels). Bottom left, FACS analysis of EEF1A2 levels in U937 cells, after 1 month of co-culture with HS5 transfected as indicated. Bottom right, histogram representing fold change in mean of fluorescence intensity relative to educated U937 maintained on HS5 transfected with siCtrl ± SEM calculated from the analysis of 3 independent experiments. Statistical analysis was performed using the nonparametric Mann-Whitney U test (***, $P<0.0005$).

**Fig. S5 (related to figure 5)**. **Effect of miR-155 on endoglin distribution, and endoglin-GIPC1 versus endoglin-syntenin interaction in surface plasmon resonance experiments. (A)** Left, FACS analysis of endoglin levels in HS5, transfected with miR-155-5p (miR-155) or control (miR-Ctrl). Cells were immunostained with the anti-human endoglin monoclonal antibody or isotype control antibody (untreated). Right, data represent the mean of the mean of fluorescence intensity ± SEM, collected from 3 independent experiments. Statistical analysis was performed using the nonparametric Mann-Whitney U test (**, $P < 0.001$). **(B)** HS5 cells were transfected with miR-155-5p (miR-155) or control (miR-Ctrl). After 48h, the media were changed and HS5 transfected cells were cultured in medium containing EV-depleted FCS (10%) for 16h. Conditioned medium was submitted to differential centrifugation and small EVs were pelleted at 100,000g. Total cell lysates and corresponding pelleted extracellular particles (small EVs) were analyzed for indicated markers. **(C)** Surface plasmon resonance illustrating the direct interaction of endoglin with GIPC1, used as positive control. BIAcore sensorgrams illustrating the binding of increasing concentrations of recombinant GIPC1 (0.5µM to 120µM) to an immobilized peptide corresponding to the last 25 C-terminal amino acids of wild-type endoglin. **(D)** Langmuir graph showing the binding (in resonance units, RU) of increasing concentrations of recombinant syntenin and GIPC1 constructs as observed at equilibrium in BIAcore. KD (apparent) was calculated from the protein concentration required to observe half of the maximal response.

**Fig. S6 (related to figure 6). Endoglin/syntenin suppression in HS5 cells (control transfection experiments). (A)** Western blots of total cell lysates from HS5 transfected with control siRNA (siCtrl) or with siRNA targeting syntenin and control (siSynt + siCtrl), endoglin and control (siEng + siCtrl), syntenin and endoglin (siSynt + siEng; 30nM) showing the effects on syntenin levels. GAPDH was used as control. **(B)** FACS analysis of endoglin surface expression on HS5 transfected with control siRNA (siCtrl, dark gray), or with siRNA targeting syntenin and control (siSynt + siCtrl, light gray), endoglin and control (siEng + siCtrl, blue), syntenin and endoglin (siSynt + siEng; red). Isotype control antibody was used as control (unstained).

**Supplementary Table S1. Antibodies and reagents**

**Supplementary table S2. Gene differentially expressed in AML-BMSC from GSE97194 microarray dataset, related to figure 1A.**

**Supplementary table S3. Gene differentially expressed in AML-BMSC depending on AML subtypes, related to figure 1B.**

**Supplementary Table S4. List of primers for RT-qPCR analysis.**

**Supplementary Table S5. FLB1-G4WT and FLB1G4-KO proteomic dataset, related to figure 2A & 2B.**
